# Glucose-dependent glycosphingolipid biosynthesis fuels CD8^+^ T cell function and tumor control

**DOI:** 10.1101/2024.10.10.617261

**Authors:** Joseph Longo, Lisa M. DeCamp, Brandon M. Oswald, Robert Teis, Alfredo Reyes-Oliveras, Michael S. Dahabieh, Abigail E. Ellis, Michael P. Vincent, Hannah Damico, Kristin L. Gallik, Shelby E. Compton, Colt D. Capan, Kelsey S. Williams, Corinne R. Esquibel, Zachary B. Madaj, Hyoungjoo Lee, Dominic G. Roy, Connie M. Krawczyk, Brian B. Haab, Ryan D. Sheldon, Russell G. Jones

## Abstract

Glucose is essential for T cell proliferation and function, yet its specific metabolic roles *in vivo* remain poorly defined. Here, we identify glycosphingolipid (GSL) biosynthesis as a key pathway fueled by glucose that enables CD8^+^ T cell expansion and cytotoxic function *in vivo*. Using ^13^C-based stable isotope tracing, we demonstrate that CD8^+^ effector T cells use glucose to synthesize uridine diphosphate-glucose (UDP-Glc), a precursor for glycogen, glycan, and GSL biosynthesis. Inhibiting GSL production by targeting the enzymes UGP2 or UGCG impairs CD8^+^ T cell expansion and cytolytic activity without affecting glucose-dependent energy production. Mechanistically, we show that glucose-dependent GSL biosynthesis is required for plasma membrane lipid raft integrity and aggregation following TCR stimulation. Moreover, UGCG-deficient CD8^+^ T cells display reduced granzyme expression and tumor control *in vivo*. Together, our data establish GSL biosynthesis as a critical metabolic fate of glucose—independent of energy production—required for CD8^+^ T cell responses *in vivo*.

## INTRODUCTION

The adaptive immune system plays a critical role in host defense against infections and cancer. T cells are central players in the adaptive immune response and respond to antigen-specific signals by activating, proliferating, and differentiating into effector T cell subsets tailored to identify and eliminate threats to the host. Reprogramming of cellular metabolism is integral to T cell responses. Upon T cell receptor (TCR) engagement and appropriate co-stimulation, naïve T (Tn) cells dramatically alter their metabolic activity as they become activated and differentiate into effector T (Teff) cells^1–3^. This metabolic reprogramming is required to meet the increased metabolic demands of cell growth, proliferation, and effector function^1–3^.

A critical step in the formation of functional Teff cells is the increased expression of glucose transporters at the plasma membrane and uptake of glucose from the extracellular environment^4–6^. T cells activated *in vitro* predominantly metabolize glucose via aerobic glycolysis and produce large amounts of lactate, a process known as the Warburg effect^7–9^. This shift towards increased glycolysis is critical, as restricting glucose availability or inhibiting glycolysis impairs T cell survival, proliferation, and cytokine production *in vitro*^4,7,10–13^. However, in addition to aerobic glycolysis, T cell activation also promotes an increase in oxidative phosphorylation (OXPHOS)^7,14^, which can be fueled in part by glucose-derived pyruvate^9,15^. Interestingly, compared to *in vitro*-activated CD8^+^ T cells, antigen-specific CD8^+^ T cells activated *in vivo* display reduced lactate production and increased rates of OXPHOS^9^. Under physiologic conditions, however, CD8^+^ T cells preferentially oxidize non-glucose carbon sources to fuel tricarboxylic acid (TCA) cycle metabolism and OXPHOS^15–17^, and instead use glucose to support anabolic growth pathways such as nucleotide^9,15^ and lipid^18^ biosynthesis. Together, these studies indicate that CD8^+^ T cells *in vivo* have distinct requirements for glucose other than energy production; however, the metabolic fates of glucose critical for supporting Teff cell responses under physiologic conditions remain poorly defined.

Here, we use *in vivo* ^13^C-glucose tracing to track metabolic fates of glucose that are critical for effector function. We demonstrate that CD8^+^ Teff cells responding to infection preferentially use glucose to synthesize the nucleotide sugar uridine diphosphate-glucose (UDP-Glc), a critical metabolic intermediate for glycogen, glycan, and glycosphingolipid (GSL) biosynthesis. We further demonstrate that inhibiting glucose-dependent UDP-Glc biosynthesis—by depleting the enzyme UDP-Glc pyrophosphorylase 2 (UGP2)—significantly impairs T cell expansion in response to infection *in vivo*. Mechanistically, we demonstrate that glucose-derived UDP-Glc is required for *de novo* GSL biosynthesis. Moreover, inhibiting GSL production—by targeting UDP-Glc ceramide glucosyltransferase (UGCG)—impairs CD8^+^ T cell expansion *in vivo* without affecting bioenergetics or energy production. This proliferative defect is associated with reduced accumulation of GSLs in the plasma membrane. GSLs are integral components of lipid rafts, microdomains at the plasma membrane that are required for optimal cell signaling. Blocking GSL biosynthesis by targeting UGCG expression impairs lipid raft aggregation following TCR stimulation. T cells with reduced glucose-dependent GSL biosynthesis display reduced granzyme expression, compromising CD8^+^ T cell cytotoxic function and tumor control *in vivo*. Together, our data highlight GSL biosynthesis as an essential metabolic fate of glucose required to maintain membrane lipid raft integrity, CD8^+^ T cell proliferation, and cytotoxic function *in vivo*.

## RESULTS

### UDP-Glc biosynthesis is a major metabolic fate of glucose in physiologically activated CD8^+^ T cells

To delineate the metabolic fates of glucose in CD8^+^ Teff cells *in vivo*, we employed a stable isotope infusion strategy developed and previously described by our group^9,19^. This approach uses CD8^+^ OT-I cells expressing a transgenic TCR specific for ovalbumin (OVA) and the CD90.1 (Thy1.1) variant of CD90, enabling us to distinguish and rapidly isolate antigen-specific T cells from hosts. Thy1.1^+^CD8^+^ OT-I cells were adoptively transferred into Thy1.2^+^ C57BL/6J hosts, which were subsequently infected with an attenuated strain of *Listeria monocytogenes* (*Lm*) expressing OVA (*Lm*-OVA) (**Figure 1A**). At the peak of infection (3 days post-infection (dpi)), either CD8^+^CD44^low^ Tn cells or antigen-specific (Thy1.1^+^) Teff cells were isolated from spleens using magnetic beads, and analyzed immediately by liquid chromatography–mass spectrometry (LC-MS) (**Figure 1A**). CD8^+^ T cells were isolated at 3 dpi to capture early metabolic changes displayed by actively proliferating Teff cells^17^.

**Figure 1:**
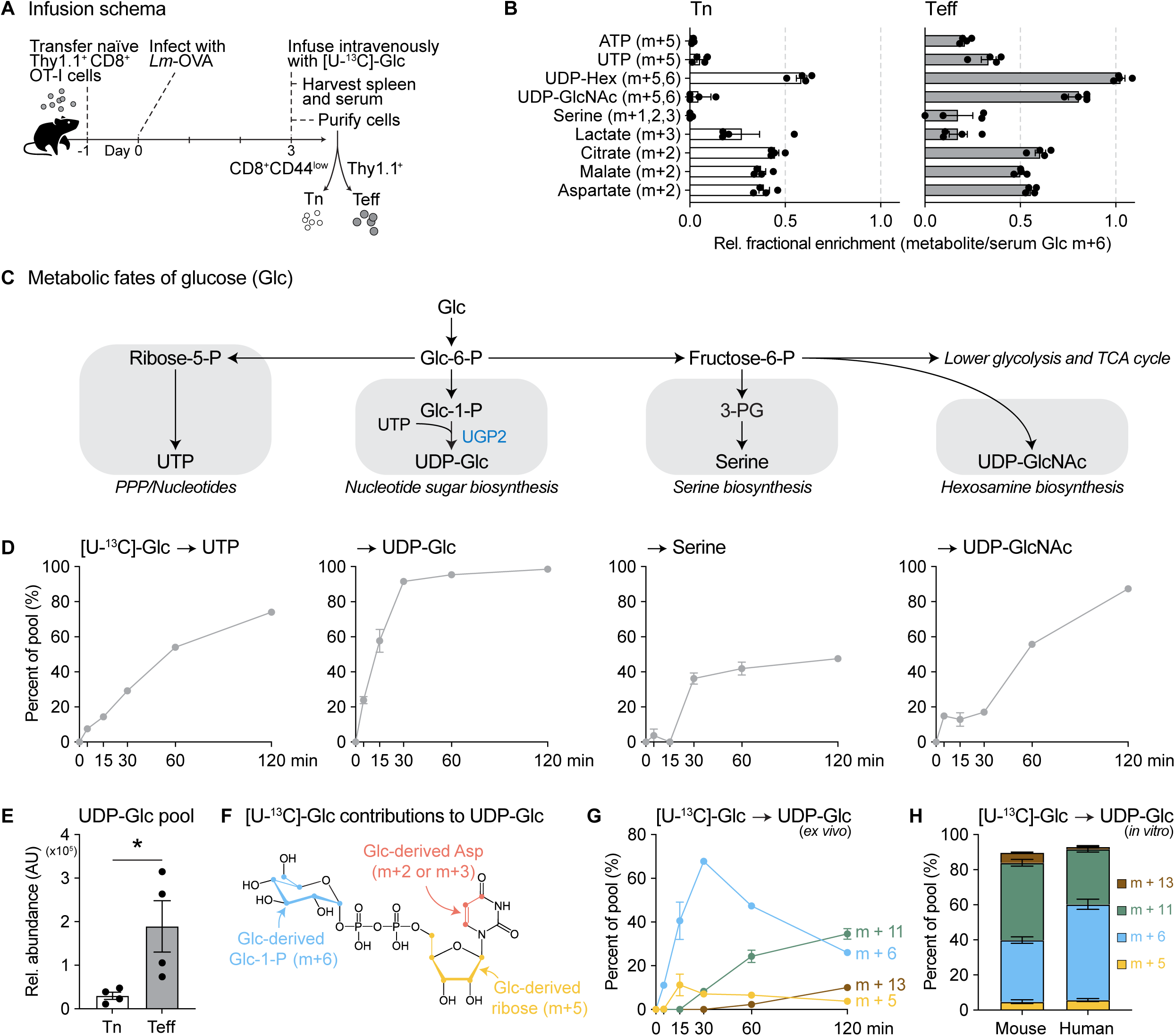
UDP-Glc biosynthesis is a major metabolic fate of glucose in physiologically activated CD8^+^ T cells. (**A**) Experimental setup for [U-^13^C]-glucose infusions in *Lm*-OVA infected mice. (**B**) Relative contribution of infused [U-^13^C]-glucose to the indicated metabolites in CD8^+^ naïve (Tn; CD8^+^CD44^low^) T cells (*left*) and *Lm*-OVA-specific effector T (Teff) cells (*right*) at 3 days post-infection (dpi). ^13^C metabolite enrichment is normalized relative to steady-state [U-^13^C]-glucose (m+6) enrichment in serum (see also Figure S1A). Data represent the mean ± SEM (n = 4 mice/group). (**C**) Schematic of major glucose-fueled metabolic pathways downstream of glucose-6-phosphate (Glc-6-P). The reaction for UGP2-dependent UDP-Glc biosynthesis is indicated. (**D**) Timecourse of [U-^13^C]-glucose incorporation into the metabolic pathways outlined in (C). CD8^+^ OT-I Teff cells isolated from *Lm*-OVA-infected mice at 3 dpi were cultured *ex vivo* for up to 2 h in VIM medium containing 5 mM [U-^13^C]-glucose. Total ^13^C enrichment (% of pool) from [U-^13^C]-glucose in UTP, UDP-Glc, serine, and UDP-GlcNAc is shown. Data represent the mean ± SEM (n = 3 biological replicates). (**E**) Relative UDP-Glc abundance in Tn and OT-I Teff cells isolated from *Lm*-OVA-infected mice at 3 dpi. Data represent the mean ± SEM (n = 4 mice/group). AU, arbitrary unit. (**F**) Schematic depicting the contribution of glucose carbon to UDP-Glc synthesis. (**G**) Mass isotopologue distribution of [U-^13^C]-glucose-derived UDP-Glc in CD8^+^ Teff cells over time. CD8^+^ OT-I Teff cells from *Lm*-OVA-infected mice as in (D) were cultured *ex vivo* for up to 2 h in VIM medium containing 5 mM [U-^13^C]-glucose. Data represent the mean ± SEM (n = 3 biological replicates). (**H**) Mass isotopologue distribution of [U-^13^C]-glucose-derived UDP-Glc in mouse and human CD8^+^ T cells. *In vitro*-activated CD8^+^ T cells were cultured for 2 h in VIM medium containing 5 mM [U-^13^C]-glucose prior to metabolite extraction. Data represent the mean ± SEM (n = 4 mice and 3 human donors).

As expected with this infusion protocol^9,19^, we achieved ∼30% enrichment of ^13^C-glucose m+6 in serum with ∼10% or less enrichment of other glucose isotopologues (**Figure S1A**). The greatest enrichment of ^13^C-glucose-derived carbon in Tn cells was observed in glycolytic (lactate) and TCA cycle-derived (citrate, malate, aspartate) metabolites, as well as uridine diphosphate (UDP)-hexose (UDP-Hex) (**Figure 1B**). Interestingly, we observed similar enrichment of ^13^C-glucose-derived carbon in the TCA cycle in activated Teff cells compared to Tn cells (**Figure 1B**). However, in contrast to Tn cells, Teff cells displayed a dramatic increase in ^13^C-glucose-dependent synthesis of metabolites generated from early glycolytic intermediates, namely nucleotides, nucleotide sugars, and serine (**Figure 1B**). In particular, nucleotide sugars, including UDP-Hex and UDP-N-acetylglucosamine (UDP-GlcNAc), were extensively labeled from glucose, indicating that nucleotide sugar biosynthesis is a major fate of glucose metabolism in CD8^+^ Teff cells.

To resolve the kinetics of glucose incorporation into nucleotide sugars relative to other glycolysis-derived metabolites (**Figure 1C**), CD8^+^ OT-I cells were activated by *Lm*-OVA infection as in **Figure 1A**, but isolated and cultured *ex vivo* in a physiologic medium (i.e., VIM^15^) containing ^13^C-glucose for up to 2 h. Cells were subjected to the same LC-MS method as in **Figure 1B**, as well as a high-pH hydrophilic interaction chromatography (HILIC) LC-MS method to better resolve different UDP-Hex species. Similar to *in vivo*-labeled CD8^+^ Teff cells, *Lm*-OVA-activated CD8^+^ Teff cells exposed to ^13^C-glucose *ex vivo* displayed ^13^C enrichment in UTP, serine, and UDP-GlcNAc, but with varying kinetics (**Figure 1D**). Strikingly, the most rapid ^13^C labeling in *Lm*-OVA-activated CD8^+^ Teff cells was seen in the nucleotide sugar UDP-glucose (UDP-Glc), with >50% of the UDP-Glc pool labeled from glucose within 15 min and ∼100% of the UDP-Glc pool labeled by 30 min (**Figure 1D**). Consistent with the rapid labeling of UDP-Glc from ^13^C-glucose, the total UDP-Glc pool was ∼6-fold greater in Teff cells compared to Tn cells at 3 dpi (**Figure 1E**). UDP-Glc-derived metabolites including UDP-galactose (UDP-Gal) and UDP-glucuronic acid (UDP-GlcA) were also rapidly labeled from ^13^C-glucose *ex vivo*, with kinetics of UDP-Gal labeling similar to that of UDP-Glc (**Figure S1B**).

UDP-Glc is synthesized from glucose-1-phosphate (Glc-1-P) and UTP, both of which can be derived from ^13^C-glucose (**Figure 1F**). A fully labeled glucose molecule ([U-^13^C]-glucose) can generate UDP-Glc m+6 from Glc-1-P, while glucose metabolized via the pentose phosphate pathway (PPP) will generate ribose m+5 and subsequently UTP m+5 (**Figures 1C** and **S1C-D**). Finally, glucose-derived aspartate can also contribute carbon to the nitrogenous base of UTP, resulting in an additional 2-3 ^13^C-labeled carbons (to generate UTP m+7 or m+8) depending on whether glucose-derived pyruvate enters the TCA cycle via pyruvate dehydrogenase (m+2) or pyruvate carboxylase (m+3) (**Figure S1D**). Isotopologue distribution analysis of UDP-Glc production in *Lm*-OVA-activated CD8^+^ Teff cells over time revealed an early increase in UDP-Glc m+6, followed by increased proportions of UDP-Glc m+11 and m+13 isotopologues by 2 h (**Figure 1G**). This labeling patten suggests that glucose-derived Glc-1-P is used first to synthesize UDP-Glc, and then m+5 UTP (labeled ribose sugar) and m+7 UTP (labeled ribose sugar and nitrogenous base) increasingly contribute to UDP-Glc synthesis over time. Importantly, the rapid synthesis of UDP-Glc in CD8^+^ Teff cells is not specific to *Lm*-OVA infection, as both murine and human CD8^+^ T cells activated *in vitro* with anti-CD3 and anti-CD28 antibodies similarly use glucose to synthesize UDP-Glc (**Figure 1H**). Collectively, these data reveal UDP-Glc biosynthesis as a major fate of glucose metabolism in CD8^+^ Teff cells, and that rapid synthesis of UDP-Glc is coordinated by sourcing carbon from multiple glucose-derived metabolic pathways in CD8^+^ Teff cells.

### UGP2 coordinates UDP-Glc biosynthesis in T cells to maintain T cell homeostasis and fuel CD8^+^ Teff responses to infection

Despite UDP-Glc being a major fate of glucose in Teff cells, the contribution of UDP-Glc metabolism to CD8^+^ T cell function is unknown. To test the importance of this pathway to T cell function *in vivo,* we generated *Ugp2*-floxed (*Ugp2*^fl/fl^) mice and crossed them with the *Cd4*-*Cre* transgenic mice to generate a mouse model with conditional deletion of *Ugp2* in mature T cells (**Figure 2A**). Baseline immunophenotyping of *Ugp2*^fl/fl^*Cd4*-*Cre* mice revealed a significant reduction in peripheral CD3^+^ T cells compared to *Cd4*-*Cre*-negative control littermates, irrespective of sex (**Figures 2B-C**). Within the CD3^+^ T cell subset, we observed a slight reduction in the percentage of CD4^+^ and CD8^+^ T cells, which translated to a 5-6-fold reduction in the total number of mature CD4^+^ and CD8^+^ T cells in the periphery (**Figures 2D-F**). Splenic B cell and NK cell numbers were similar between genotypes (**Figure S2A**).

**Figure 2:**
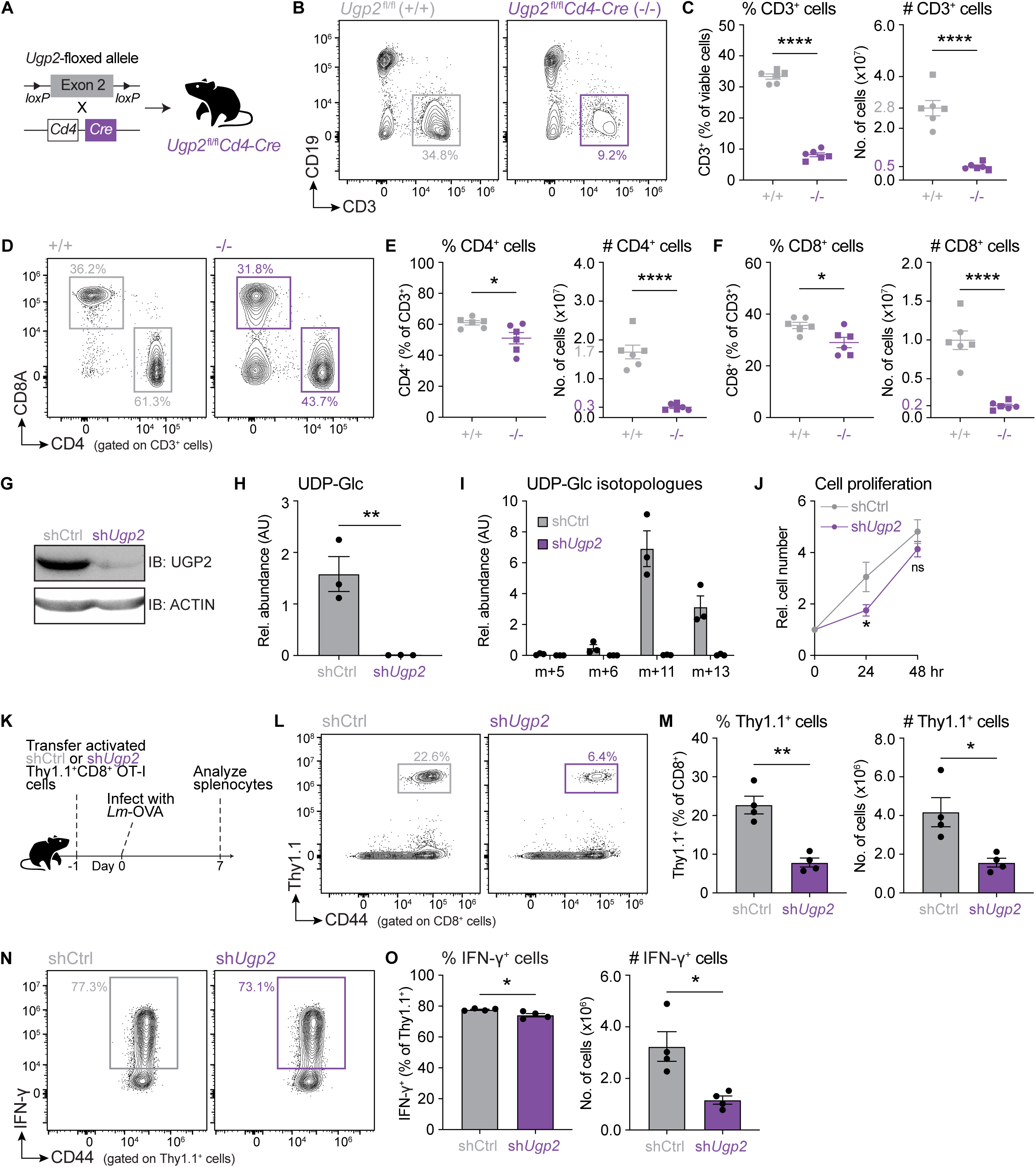
UGP2 coordinates UDP-Glc biosynthesis in T cells to maintain T cell homeostasis and fuel CD8^+^ Teff cell responses to infection. (**A**) Targeting strategy for T cell-specific deletion of *Ugp2* in mice (*Ugp2*^fl/fl^*Cd4*-*Cre* strain). (**B**) Representative flow cytometry plots for CD19 versus CD3 expression from splenocytes isolated from control *Ugp2*^fl/fl^ (+/+) and knockout *Ugp2*^fl/fl^*Cd4*-*Cre* (−/−) mice. (**C**) Percentage (*left*) and total number (*right*) of CD3^+^ T cells in the spleen of *Ugp2*^fl/fl^ (+/+) and *Ugp2*^fl/fl^*Cd4*-*Cre* (−/−) mice. Data represent the mean ± SEM (n = 6 mice/group; circle = female, square = male). (**D**) Representative flow cytometry plots for CD8A versus CD4 expression (gated on CD3^+^ cells) from splenocytes isolated from *Ugp2*^fl/fl^ (+/+) and *Ugp2*^fl/fl^*Cd4*-*Cre* (−/−) mice. (**E-F**) Peripheral T cell populations in T cell-specific UGP2-deficient mice. Percentage (*left*) and total number (*right*) of CD4^+^ T cells (E) and CD8^+^ T cells (F) in the spleen of control *Ugp2*^fl/fl^ (+/+) and knockout *Ugp2*^fl/fl^*Cd4*-*Cre* (−/−) mice. Data represent the mean ± SEM (n = 6 mice/group; circle = female, square = male). (**G**) Immunoblot of UGP2 protein expression in control shRNA (shCtrl) and sh*Ugp2*-expressing CD8^+^ T cells. ACTIN protein expression is shown as a loading control. (**H**) Relative abundance of UDP-Glc in shCtrl and sh*Ugp2*-expressing CD8^+^ T cells cultured *in vitro*. Data represent the mean ± SEM (n = 3). AU, arbitrary unit. (**I**) Relative abundance of the indicated [U-^13^C]-glucose-derived UDP-Glc mass isotopologues in shCtrl and sh*Ugp2*-expressing CD8^+^ T cells after 24 h of culture in VIM medium containing 5 mM [U-^13^C]-glucose. Data represent the mean ± SEM (n = 3). AU, arbitrary unit. (**J**) Relative cell number over time for activated shCtrl and sh*Ugp2*-expressing CD8^+^ T cells cultured *in vitro* in IMDM containing 50 U/mL IL-2. Data represent the mean ± SEM (n = 3). (**K-M**) Expansion of UGP2-depleted CD8^+^ OT-I cells *in vivo* in response to *Lm*-OVA infection. (K) Experimental setup for adoptive transfer of control (shCtrl) or sh*Ugp2*-expressing Thy1.1^+^CD8^+^ OT-I cells followed by *Lm*-OVA infection. (**L**) Representative flow cytometry plots showing the abundance of antigen-specific (Thy1.1^+^) control and sh*Ugp2*-expressing CD8^+^ OT-I cells in the spleen of *Lm*-OVA-infected mice at 7 days post-infection (dpi). (**M**) Percentage (*left*) and total number (*right*) of antigen-specific (Thy1.1^+^) control and sh*Ugp2*-expressing CD8^+^ OT-I cells in the spleen of *Lm*-OVA-infected mice at 7 dpi. Data represent the mean ± SEM (n = 4 mice/group). (**N-O**) Cytokine response of UGP2-depleted CD8^+^ OT-I cells *ex vivo*. (N) Representative flow cytometry plots showing the percentage of IFN-γ-producing Thy1.1^+^ control (shCtrl) and sh*Ugp2*-expressing CD8^+^ OT-I cells in the spleen of *Lm*-OVA-infected mice at 7 dpi after *ex vivo* re-stimulation with OVA peptide. (**O**) Percentage (*left*) and total number (*right*) of IFN-γ-producing Thy1.1^+^ control and sh*Ugp2*-expressing CD8^+^ OT-I cells in the spleen of *Lm*-OVA-infected mice at 7 dpi after *ex vivo* re-stimulation with OVA peptide. Data represent the mean ± SEM (n = 4 mice/group).

To identify whether the decrease in peripheral T cells was due to impaired T cell development, we profiled T cell subsets in the thymus. Interestingly, we observed no difference in the percentage or number of CD4 and CD8 double-negative, double-positive, or single-positive T cells in the thymus (**Figure S2B-C**). Taken together, these data suggest that UGP2 is required for T cell homeostasis outside the thymus. Notably, mature CD3^+^ T cells isolated from the spleen of *Ugp2*^fl/fl^*Cd4*-*Cre* mice retained UGP2 protein expression (**Figure S2D**) and the abundance of UDP-Glc in T cells from *Ugp2*^fl/fl^*Cd4*-*Cre* mice was similar to control cells (**Figure S2E**). These data suggest that UPG2 is essential for peripheral T cell homeostasis and that the remaining mature T cells in *Ugp2*^fl/fl^*Cd4*-*Cre* mice escaped *Ugp2* gene deletion.

Given that UGP2 appears to be essential for mature peripheral T cell homeostasis, we silenced *Ugp2* in purified, *in vitro*-activated CD8^+^ T cells using a short hairpin RNA (shRNA) as an alternative strategy. Compared to a control shRNA targeting firefly luciferase (shCtrl), expression of a *Ugp2*-targeting shRNA (sh*Ugp2*) reduced UGP2 protein expression in CD8^+^ T cells (**Figure 2G**) and ablated both total and ^13^C-glucose-derived UDP-Glc pools (**Figures 2H-I**). Interestingly, the distribution of ^13^C-glucose-derived UDP-Glc isotopologues in sh*Ugp2*-expressing CD8^+^ T cells was similar to control cells (**Figure S2F**), indicating that this strategy reduced the overall rate of UDP-Glc synthesis rather than changing its source of production.

We next examined the impact of silencing UGP2 on CD8^+^ T cell function. Knockdown of *Ugp2* in activated CD8^+^ T cells did not dramatically compromise cell proliferation *in vitro* (**Figure 2J**), nor did it alter markers of T cell activation (**Figure S2G**). However, reducing UGP2-dependent glucose metabolism had a dramatic impact on Teff cell expansion *in vivo*. We adoptively transferred shCtrl- or sh*Ugp2*-expressing Thy1.1^+^CD8^+^ OT-I cells into Thy1.2^+^ hosts, and then infected the mice with *Lm*-OVA the following day (**Figure 2K**)^9,15^. We observed a significant reduction in the percentage of antigen-specific (Thy1.1^+^) sh*Ugp2*-expressing CD8^+^ Teff cells at the peak of infection (7 dpi), which translated to a ∼2.5-fold decrease in the total number of antigen-specific cells compared to controls (**Figures 2L-M**). Despite this reduction in antigen-specific CD8^+^ T cell expansion, UGP2-depleted cells displayed no major defects in their ability to produce cytokines or differentiate into effector or memory precursor subsets (**Figures 2M-N** and **S2H-K**). Thus, our data indicate that UGP2, despite having minimal impact on CD8^+^ T cell proliferation or survival *in vitro*, is critical for both T cell homeostasis and expansion in response to infection *in vivo*.

### UDP-Glc fuels glycogen and glycan biosynthesis in CD8^+^ T cells

We next sought out to identify the metabolic pathway(s) downstream of UDP-Glc biosynthesis required for CD8^+^ Teff cell responses *in vivo*. UDP-Glc is a central intermediate for the synthesis of several metabolites, including glycogen, UDP-Gal, glycans, glycosaminoglycans, and GSLs (**Figure 3A**). Glycogen is a glucose polysaccharide that serves as a form of carbon storage in animals. A previous study reported that glycogen pools are elevated in CD8^+^ memory T (Tmem) cells compared to Tn and Teff cells, and that glycogen storage is required for the formation of T cell memory^20^. Concordantly, impairing glycogen mobilization has been shown to impair CD8^+^ Tmem cell responses *in vivo*, while having no effect on primary Teff cell responses following *Lm*-OVA infection^21,22^. We observed a significant decrease in glycogen pools in sh*Ugp2*-expressing CD8^+^ T cells compared to controls (**Figure 3B**); however, since we also found that UGP2 depletion significantly impairs CD8^+^ Teff cell expansion following *Lm*-OVA infection (**Figures 2L-M**), we decided to quantify other UDP-Glc-derived metabolites in sh*Ugp2*-expressing CD8^+^ T cells.

**Figure 3:**
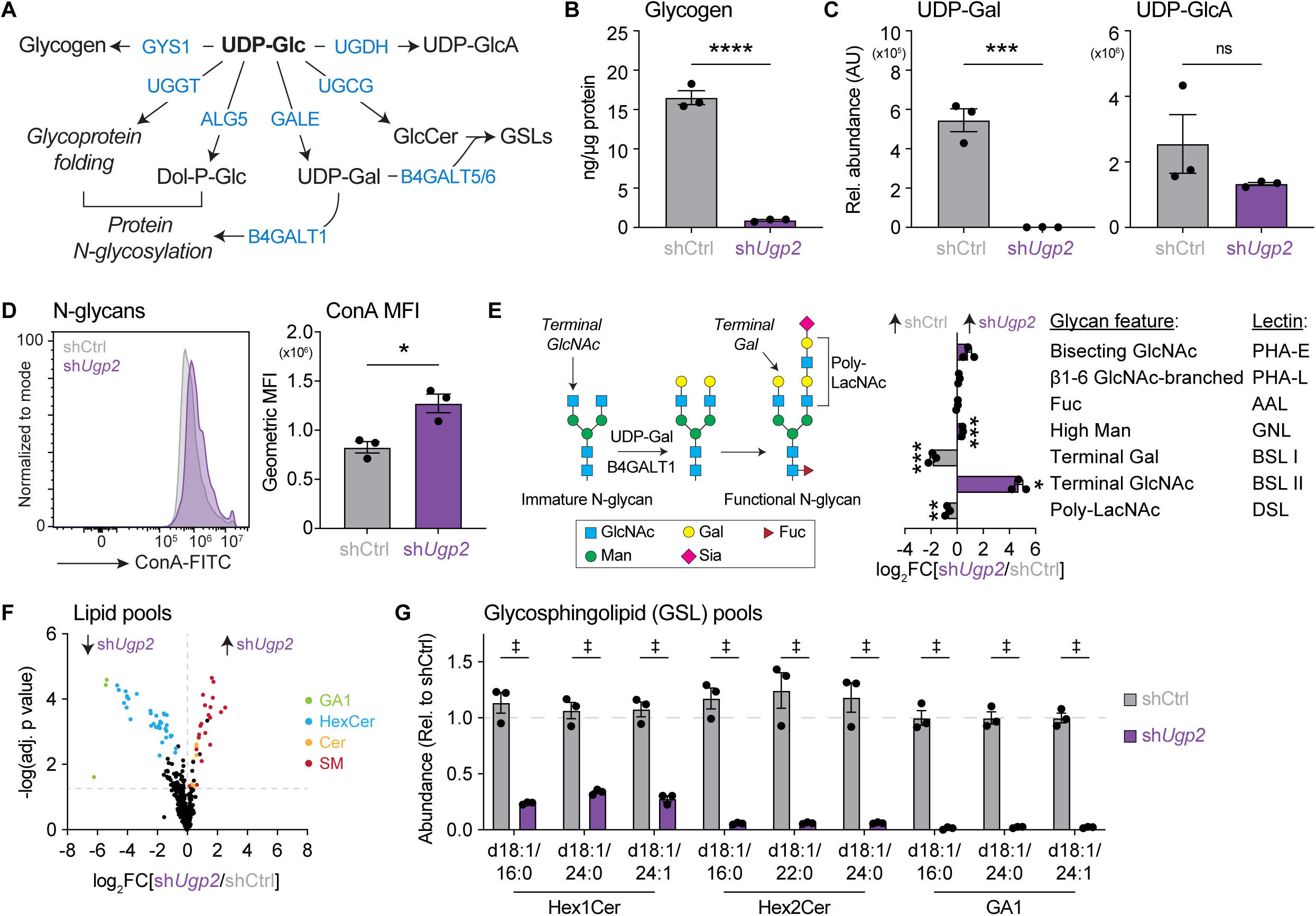
UDP-Glc fuels glycogen biosynthesis and UDP-Gal-dependent metabolic processes in CD8^+^ T cells. (**A**) Schematic showing metabolic pathways fueled by UDP-Glc. Metabolic intermediates are shown in black and enzymes are shown in blue. (**B**) Total glycogen abundance in *in vitro*-activated shCtrl- (control) and sh*Ugp2*-expressing CD8^+^ T cells. Data represent the mean ± SEM (n = 3). (**C**) Relative abundance of UDP-Gal (*left*) and UDP-GlcA (*right*) levels in *in vitro*-activated control and sh*Ugp2*-expressing CD8^+^ T cells. Data represent the mean ± SEM (n = 3). AU, arbitrary unit. (**D**) N-glycan expression on the surface of activated control and sh*Ugp2*-expressing CD8^+^ T cells. Representative histogram (*left*) and quantification of geometric mean fluorescence intensity (MFI; *right*) of ConA on control and sh*Ugp2*-expressing CD8^+^ T cells. Data represent the mean ± SEM (n = 3). (**E**) Glycan profiling of *in vitro*-activated control and sh*Ugp2*-expressing CD8^+^ T cells via lectin binding. *Left*, Schematic of N-glycan extension. Sugar structures are highlighted in the legend (GlcNAc, N-acetylglucosamine; Gal, galactose; Fuc, fucose; Man, mannose; Sia, sialic acid). *Right*, geometric MFI of the indicated glycan features on the surface of CD8^+^ T cells and the lectins that bind them. Data are plotted as the log_2_ fold change (log_2_FC) of sh*Ugp2*/shCtrl. Data represent the mean ± SEM (n = 3). (**F**) Volcano plot of lipid levels in *in vitro*-activated control and sh*Ugp2*-expressing CD8^+^ T cells. Specific lipid species are indicated (GA1, gangliosides; HexCer, hexosylceramides; Cer, ceramides; SM, sphingomyelins). Data are plotted as the log_2_ fold change (log_2_FC) of sh*Ugp2*/shCtrl and represent the average of 3 biological replicates. The horizontal line indicates the false discovery rate (FDR)-adjusted p value cutoff of 0.05 and the vertical line indicates a log_2_FC of 0. (**G**) Relative abundance of the indicated GSLs in *in vitro*-activated sh*Ugp2*-expressing CD8^+^ T cells relative to the mean abundance in shCtrl-expressing cells. Data represent the mean ± SEM (n = 3). Statistical significance was determined using multiple t-tests and an FDR correction for multiple comparisons. ‡, q < 0.01.

Metabolomics analysis of control and sh*Ugp2*-expressing CD8^+^ T cells revealed a significant decrease in the pool of UDP-Gal in UGP2-depleted cells, while the pool of UDP-GlcA, a precursor for glycosaminoglycan biosynthesis, was similar between control and sh*Ugp2*-expressing cells (**Figure 3C**). UDP-Gal serves as a substrate for β-1,4-galactosyltransferases (B4GALTs), which catalyze the synthesis of glycoproteins and glycolipids^23,24^. While evidence for functional redundancies between B4GALT enzymes exists^23,24^, B4GALT1 primarily catalyzes the transfer of Gal to terminal GlcNAc acceptors on complex-type N-glycans^25^, whereas B4GALT5/6 catalyze UDP-Gal-dependent synthesis of lactosylceramide (LacCer) and GSLs^26,27^ (**Figure 3A**). In addition to UDP-Gal-dependent galactosylation of glycoproteins, UDP-Glc can also regulate glycoprotein folding and N-glycan initiation at the endoplasmic reticulum^28,29^. To interrogate whether silencing *Ugp2* in CD8^+^ T cells impacted protein N-glycosylation, we probed cells with the mannose-binding lectin Concanavalin A (ConA), which is commonly used to detect N-glycans at the cell surface^30^. Surprisingly, we observed a slight, statistically significant increase in the mean fluorescence intensity (MFI) of ConA on sh*Ugp2*-expressing CD8^+^ T cells (**Figure 3D**), indicating that UGP2-depleted cells either have a greater number of N-glycans on their surface or an altered N-glycan composition that favors ConA binding.

N-glycans undergo maturation in the Golgi apparatus, where B4GALT1 transfers Gal from UDP-Gal to terminal GlcNAc acceptors to form N-acetyllactosamine (LacNAc) and initiate N-glycan extension^31^ (**Figure 3E**). To further interrogate the phenotype of N-glycans in sh*Ugp2*-expressing CD8^+^ T cells, we probed cells with a panel of lectins that recognize different N-glycan features. Interestingly, sh*Ugp2*-expressing CD8^+^ T cells displayed a significant increase in terminal GlcNAc and a decrease in terminal Gal as determined by binding of *Bandeiraea simplicifolia* lectin (BSL) II and BSL I, respectively (**Figure 3E**). Moreover, we observed a decrease in poly-LacNAc features in sh*Ugp2*-expressing CD8^+^ T cells, as evidenced by reduced MFI of *Datura stramonium* lectin (DSL) (**Figure 3E**). Taken together, these data indicate that blocking glucose-dependent UDP-Gal production (via UGP2 depletion) impairs N-glycan extension.

Consistent with the increase in ConA binding to UGP2-depleted CD8^+^ T cells (**Figure 3D**), we also observed a slight but significant increase in binding of another mannose-binding lectin, *Galanthus nivalis* lectin (GNL), in UGP2-depleted cells (**Figure 3E**). Without proper N-glycan extension, the mannose residues in the core of the truncated N-glycans (**Figure 3E**) may become more exposed to mannose-binding lectins like ConA and GNL. Indeed, these lectins typically exhibit lower affinity for core mannose residues when additional carbohydrate modifications, such as branch extensions, are present^32,33^. Hence, silencing *Ugp2* and disrupting glucose-dependent UDP-Gal biosynthesis remodels the composition of the glycocalyx—glycan coating on the cell surface—of CD8^+^ T cells.

### UGP2 is required for GSL biosynthesis in CD8^+^ T cells

In addition to their roles in N-glycosylation, UDP-Glc and UDP-Gal are also substrates for GSL biosynthesis (**Figure 3A**). The enzyme UGCG catalyzes the first step in GSL biosynthesis by transferring Glc from UDP-Glc to a ceramide molecule to produce glucosylceramide (GlcCer) (**Figure 3A**). B4GALT5 then transfers Gal from UDP-Gal to GlcCer to produce LacCer, which goes on to produce a wide array of GSLs including gangliosides^26,27,34^. To determine how depleting UGP2 impacted GSL synthesis, we conducted lipidomics analysis of control and sh*Ugp2*-expressing CD8^+^ T cells. The most robust change we observed was a decrease in numerous hexosylceramide (HexCer) and GA1 ganglioside species in sh*Ugp2*-expressing T cells (**Figures 3F-G** and **S3A-B**). This decrease in HexCer and ganglioside species was accompanied by a slight increase in ceramides and ceramide-derived sphingomyelins (**Figure 3F**), suggesting a re-routing of ceramides from HexCer biosynthesis towards alternative lipid metabolic pathways when UGP2 is depleted. Notably, depletion of UDP-Glc and GSL pools in UGP2-depleted CD8^+^ T cells had little-to-no impact on the abundance or ^13^C labeling of other glucose-derived metabolites linked to T cell proliferation, including adenosine triphosphate (ATP), serine, UDP-GlcNAc, and citrate (**Figure S3C-D**). Collectively, these data indicate that UGP2 regulates the production of multiple UDP-Gal-dependent metabolic processes in CD8^+^ T cells, without impacting other glucose-dependent biosynthetic and bioenergetic pathways.

### CD8^+^ Teff cells direct glucose to fuel GSL biosynthesis in vivo

To identify how UDP-Gal-dependent metabolic pathways change with T cell activation, we mined previously published proteomics data from CD8^+^ Tn and Teff cells isolated from *Lm*-OVA-infected mice^9^. Relative to Tn cells, CD8^+^ Teff cells at 3 dpi displayed increased expression of the enzyme UDP-Gal-4-epimerase (GALE) (**Figure 4A**), which catalyzes the conversion of UDP-Glc to UDP-Gal (**Figures 3A**). This upregulation of GALE expression with T cell activation is consistent with the robust glucose-dependent synthesis of UDP-Gal observed in activated CD8^+^ T cells (**Figure S1B**). Teff cells also displayed higher expression of UGCG and B4GALT5 relative to Tn cells, while the expression of B4GALT1, which directs UDP-Gal to protein N-glycosylation, was similar between Teff and Tn cells (**Figure 4A**). Taken together, these data indicate metabolic reprogramming in activated CD8^+^ T cells directs glucose towards increased GSL biosynthesis *in vivo*.

**Figure 4:**
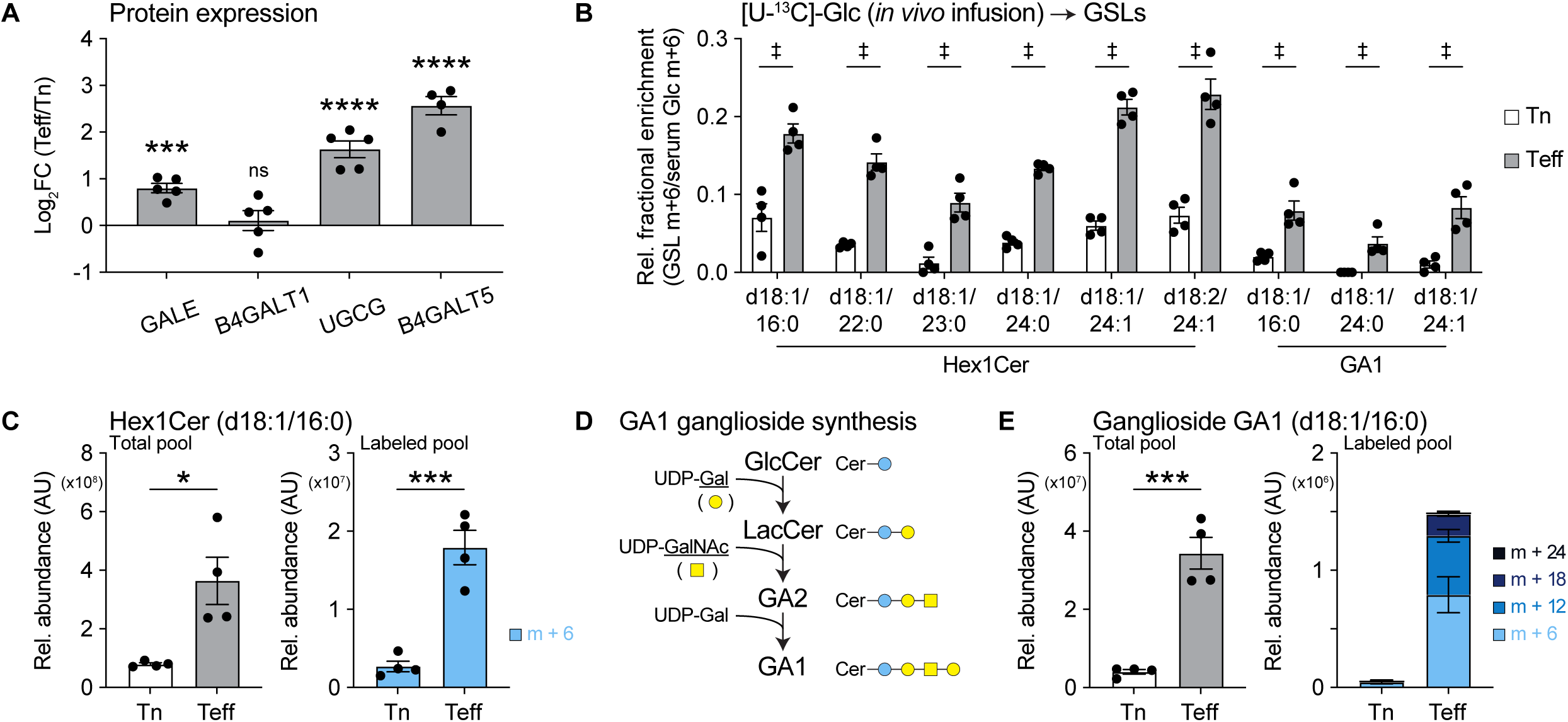
CD8^+^ T cells direct glucose to fuel GSL biosynthesis *in vivo*. (**A**) Protein levels of GSL biosynthesis pathway and galactosyltransferase enzymes in activated CD8^+^ T cells *in vivo*. Data are expressed as the log_2_ fold change (log_2_FC) in protein levels in antigen-specific CD8^+^ OT-I T effector (Teff) cells relative to CD8^+^ naïve T (Tn) cells isolated from *Lm*-OVA-infected mice at 3 days post-infection (dpi). Data represent the mean ± SEM (n = 4-5 mice/group). Data were mined from Ma *et al.*^9^. (**B**) Relative contribution of [U-^13^C]-glucose to the indicated GSL species (m+6) in CD8^+^ naïve (Tn; CD8^+^CD44^low^) T cells and *Lm*-OVA-specific Teff cells following [U-^13^C]-glucose infusion at 3 dpi. Mice were infused with [U-^13^C]-glucose *in vivo* at 3 dpi as described in Figure 1A. ^13^C metabolite enrichment is normalized relative to steady-state [U-^13^C]-glucose (m+6) enrichment in serum (see also Figure S1A). Data represent the mean ± SEM (n = 4 mice/group). Statistical significance was determined using multiple t-tests and a false discovery rate correction for multiple comparisons. ‡, q < 0.01. (**C**) Relative abundance of total (*left*) and m+6-labeled (*right*) Hex1Cer (d18:1/16:0) in Tn and Teff cells from *Lm*-OVA-infected mice infused with [U-^13^C]-glucose *in vivo* at 3 dpi. Data represent the mean ± SEM (n = 4 mice/group). AU, arbitrary unit. (**D**) Schematic of GA1 ganglioside biosynthesis from GlcCer. (**E**) Relative abundance of total (*left*) and m+6-, m+12-, m+18-, and m+24-labeled (*right*) GA1 (d18:1/16:0) ganglioside in Tn and Teff cells from *Lm*-OVA-infected mice infused with [U-^13^C]-glucose *in vivo* at 3 dpi. Data represent the mean ± SEM (n = 4 mice/group). AU, arbitrary unit.

To directly test whether CD8^+^ Teff cells actively direct glucose into GSL biosynthesis *in vivo*, we repeated the ^13^C-glucose *in vivo* infusion experiment in *Lm*-OVA-infected mice as described in **Figure 1A**, and quantified the abundance and ^13^C labeling of GSLs in purified Tn and Teff cells by LC-MS. The overall contribution of ^13^C-glucose to HexCer and GA1 ganglioside synthesis was significantly increased in CD8^+^ Teff cells at 3 dpi compared to Tn cells (**Figure 4B**). Strikingly, we observed a ∼20% relative enrichment of m+6-labeled HexCer species (i.e., d18:1/16:0, d18:1/24:1, d18:2/24:1) in CD8^+^ Teff cells after only 2 h of ^13^C-glucose infusion, indicating rapid turnover of these lipid pools. Teff cells had higher amounts of both total and ^13^C-glucose-derived HexCer and GA1 ganglioside pools compared to Tn cells, further supporting the upregulation of GSL biosynthesis in physiologically activated CD8^+^ T cells *in vivo* (**Figures 4C-E**). GA1 gangliosides are synthesized via the sequential addition of carbohydrates to GlcCer to produce a molecule comprised of a ceramide and four-sugar chain (**Figure 4D**). After 2 h of ^13^C-glucose infusion, CD8^+^ Teff cells had predominantly m+6-labeled GA1 ganglioside (one labeled sugar); however, m+12- and m+18-labeled GA1 gangliosides were also observed, suggesting that glucose contributes to the synthesis of multiple nucleotide sugars used to synthesize GA1 gangliosides (**Figure 4E**). Notably, proliferating human CD8^+^ T cells also directed glucose into GSL biosynthesis; however, GM3 was the predominant ganglioside species detected in human T cells (**Figure S4**), suggesting some level of species-dependent regulation of this metabolic pathway. Taken together, these data not only reveal that CD8^+^ Teff cells use glucose to synthesize GSLs *in vivo*, but that—due to the high turnover rate of these lipids—Teff cells actively use glucose to maintain synthesis of these lipids.

### Glucose-dependent GSL biosynthesis is essential for T cell expansion in vivo

To determine the contribution of glucose-dependent GSL biosynthesis to CD8^+^ Teff cell responses *in vivo*, we targeted the enzyme UGCG, which catalyzes the first glycosylation step in GSL biosynthesis (**Figure 3A**). We first took a pharmacological approach to inhibit UGCG enzymatic activity using the drug eliglustat^35^ (**Figure 5A**). Treating CD8^+^ T cells with a range of eliglustat concentrations (0-25 μM) during *in vitro* anti-CD3 and anti-CD28 antibody stimulation resulted in a dose-dependent decrease in cell proliferation (**Figures 5B**). At 4 μM (IC_50_ for proliferation), eliglustat significantly decreased the abundance of both HexCer and GA1 ganglioside pools (**Figure 5C**). Similarly, treating activated CD8^+^ T cells with eliglustat reduced cell proliferation in a concentration-dependent manner, with no effect on cell viability at concentrations ≤ 4 μM (**Figures S5A-B**).

**Figure 5:**
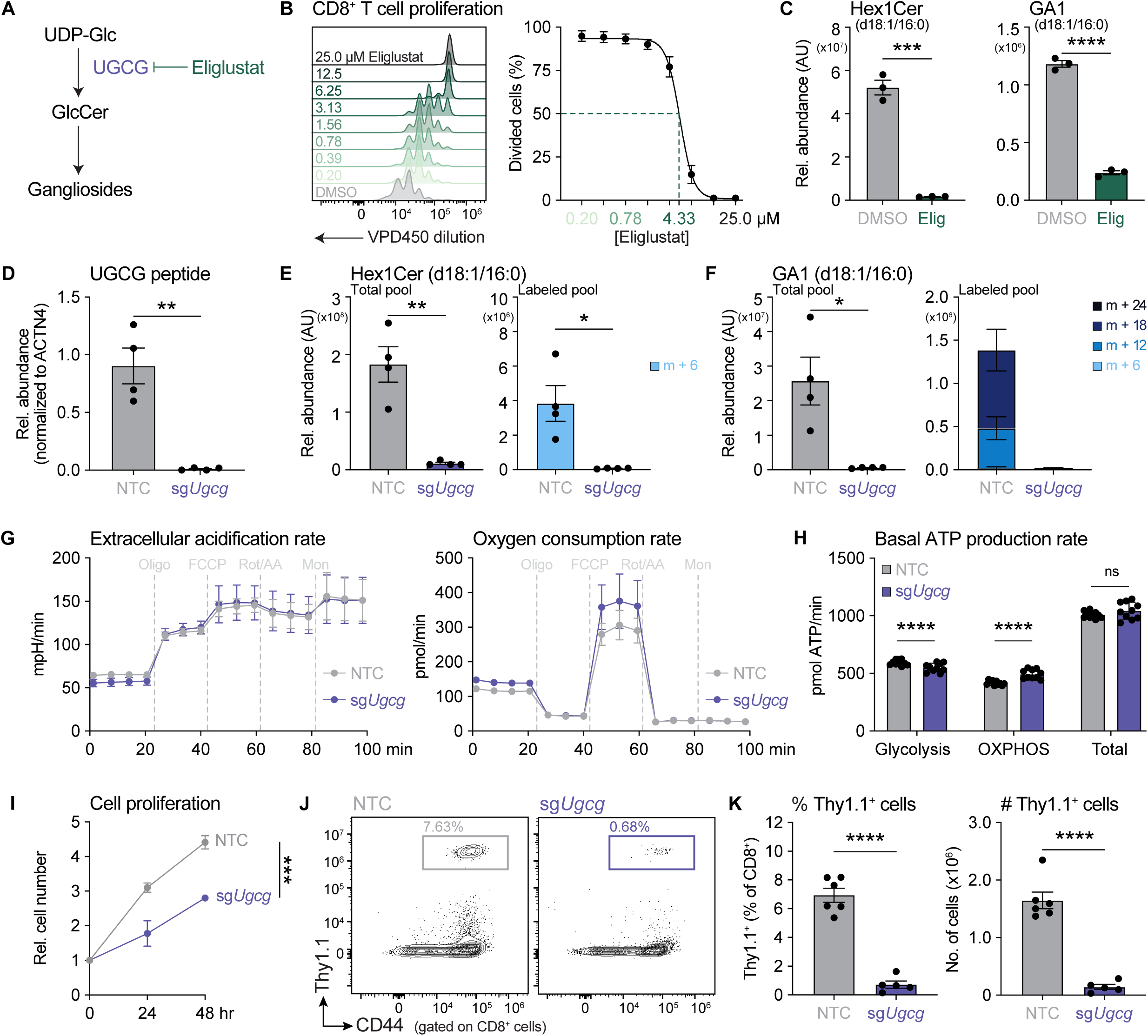
Glucose-dependent GSL biosynthesis is essential for CD8^+^ T cell expansion *in vivo.* (**A**) Schematic of the GSL biosynthesis pathway. Eliglustat inhibits GlcCer production by inhibiting the enzyme UGCG. (**B**) Proliferation of CD8^+^ T cells as measured by VPD450 dilution after 72 h of activation *in vitro* with anti-CD3 and anti-CD28 antibodies in the presence of eliglustat (0-25 μM). Representative histograms of VPD450 staining (*left*) and quantification of the percentage of divided cells (*right*) are shown. The dashed line indicates the EC_50_ of eliglustat (4.33 μM) for inhibiting proliferation. Data represent the mean ± SEM (n = 3). (**C**) Relative abundance of Hex1Cer (d18:1/16:0) (*left*) and GA1 (d18:1/16:0) ganglioside (*right*) in CD8^+^ T cells treated with DMSO or 4 μM eliglustat for 24 h. Data represent the mean ± SEM (n = 3). AU, arbitrary unit. (**D**) Relative UGCG protein abundance (normalized to ACTN4 abundance) in CD8^+^ T cells as determined by LC-MS-based proteomics. Activated CD8^+^ T cells were modified using CRISPR/Cas9 gene editing with either a non-targeting control (NTC) single guide RNA (sgRNA) or a sgRNA targeting *Ugcg* (sg*Ugcg*). Data represent the mean ± SEM (n = 4). (**E**) Relative abundance of total (*left*) and m+6-labeled (*right*) Hex1Cer (d18:1/16:0) in NTC- and sg*Ugcg*-modified CD8^+^ T cells after 24 h of culture in VIM medium containing 5 mM [U-^13^C]-glucose. Data represent the mean ± SEM (n = 4). AU, arbitrary unit. (**F**) Relative abundance of total (*left*) and m+6-, m+12-, m+18-, and m+24-labeled (*right*) GA1 (d18:1/16:0) ganglioside in NTC- and sg*Ugcg*-modified CD8^+^ T cells after 24 h of culture in VIM medium containing 5 mM [U-^13^C]-glucose. Data represent the mean ± SEM (n = 4). AU, arbitrary unit. (**G-H**) Bioenergetic profile of NTC- and sg*Ugcg*-modified CD8^+^ T cells. (G) Graphs depict the extracellular acidification rate (*left*) and oxygen consumption rate (*right*) of edited CD8^+^ T cells over time. Oligomycin (Oligo), FCCP, rotenone and antimycin A (Rot/AA), and monensin (Mon) were added to cells where indicated. (**H**) Basal ATP production rates from glycolysis, OXPHOS, and glycolysis + OXPHOS (total) in NTC- and sg*Ugcg*-modified CD8^+^ T cells. Data represent the mean ± SD (n = 10/group). (**I**) Relative cell number over time for NTC- and sg*Ugcg*-modified CD8^+^ T cells cultured *in vitro* in IMDM containing 50 U/mL IL-2. Data represent the mean ± SEM (n = 4). Statistical significance was determined using a two-way ANOVA. (**J-K**) Expansion of sg*Ugcg*-modified CD8^+^ T cells *in vivo* in response to *Lm*-OVA infection. NTC- or sg*Ugcg*-modified Thy1.1^+^CD8^+^ OT-I cells were transferred into congenic hosts, the mice were infected with *Lm*-OVA, and T cell responses in the spleen of *Lm*-OVA-infected mice were analyzed at 7 days-post infection (dpi; as in Figure 2K). (J) Representative flow cytometry plots showing the abundance of antigen-specific (Thy1.1^+^) NTC- and sg*Ugcg*-modified CD8^+^ OT-I cells in the spleen at 7 dpi. (**K**) Percentage (*left*) and total number (*right*) of antigen-specific (Thy1.1^+^) NTC- and sg*Ugcg*-modified Thy1.1^+^CD8^+^ OT-I cells from (J). Data represent the mean ± SEM (n = 5-6 mice/group).

We validated these findings using a single guide RNA (sgRNA) targeting *Ugcg*, which ablated UGCG protein expression (**Figures 5D** and **S5C**) and significantly reduced both GSL abundance and ^13^C-glucose-dependent GSL biosynthesis in CD8^+^ T cells (**Figures 5E-F**). Importantly, deletion of *Ugcg* did not significantly affect glycolysis or total ATP production, as evidenced by similar extracellular acidification and oxygen consumption rates (**Figure 5G**) and basal ATP production rates (**Figure 5H**) between control and UGCG-deficient CD8^+^ T cells. While we observed a slight decrease in basal ATP production from glycolysis in UGCG-deficient CD8^+^ T cells (**Figure 5H**), total ATP production was preserved due to a compensatory increase in ATP production from OXPHOS (**Figures 5G-H**). Furthermore, unlike with *Ugp2* silencing in CD8^+^ T cells, both glycogen pools and UDP-Gal-dependent N-glycan extension were unaffected by *Ugcg* deletion (**Figure S5D-E**). Despite having no effects on ATP production, glycogen levels, or UDP-Gal-dependent N-glycan extension, deleting *Ugcg* reduced the proliferation of CD8^+^ T cells *in vitro* (**Figure 5I**), similar to our observations with eliglustat treatment (**Figure 5B**). Moreover, similar to UGP2 depletion (**Figures 2L-M**), *Ugcg* deletion significantly reduced the number of antigen-specific CD8^+^ Teff cells *in vivo* at 7 dpi following *Lm*-OVA infection (**Figures 5J-K**), without compromising cytokine production (**Figures S5F-H**) or cell differentiation (**Figures S5I-J**). Together, these data indicate that GSL biosynthesis—regulated by the enzyme UGCG—is the key fate of glucose downstream of UGP2-mediated UDP-Glc production that is critical for CD8^+^ Teff cell expansion *in vivo*.

### GSL biosynthesis is required to maintain lipid rafts at the plasma membrane

To determine the downstream consequences of *Ugcg* deletion in CD8^+^ T cells, we performed global proteomic analysis of control and UGCG-deficient CD8^+^ T cells. We identified 37 proteins that were significantly upregulated in UGCG-deficient CD8^+^ T cells and 59 proteins that were significantly downregulated (**Figure 6A** and **Table S1**). Gene set enrichment analysis (GSEA) revealed several pathways influenced by UGCG expression (**Figures 6B** and **S6**, and **Table S2**). Notably, pathways related to TCR signal transduction and immune synapse formation, including “positive regulation of leukocyte degranulation” and “actin filament bundle assembly”, were downregulated in UGCG-deficient CD8^+^ T cells (**Figure 6B** and **Table S2**). In particular, protein expression of several granzyme family members (GZME, GZMK, GZMA) and perforin (PRF1) were significantly decreased in UGCG-deficient CD8^+^ T cells, as well as proteins implicated in actin remodeling and intracellular trafficking (ARL6, SCIN, SWAP70) (**Figure 6A**).

**Figure 6:**
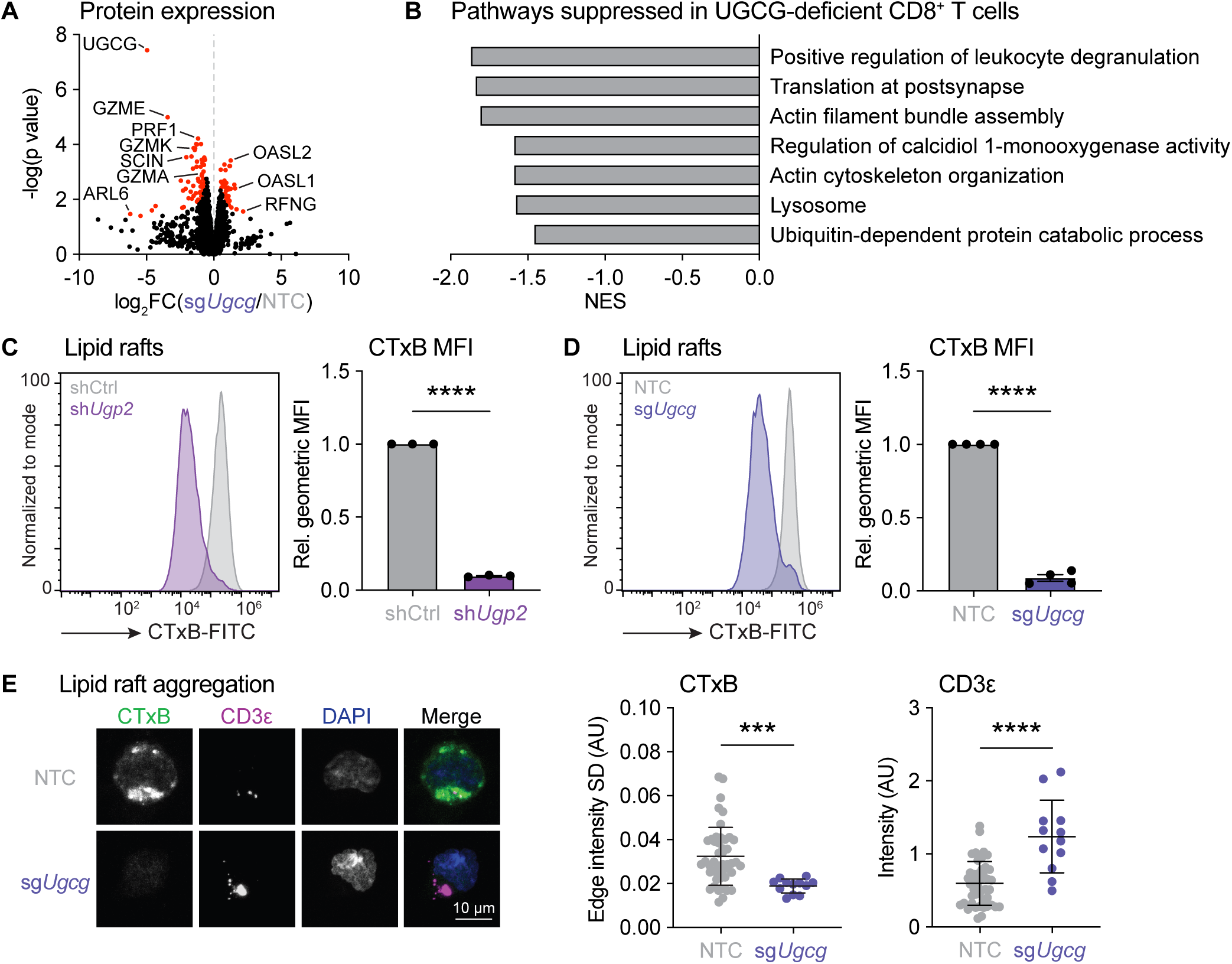
GSL biosynthesis is required to maintain lipid rafts at the plasma membrane. (**A**) Global proteomic analysis of control (NTC) and UGCG-deficient (sg*Ugcg*) CD8^+^ T cells. Data represent the average of 4 biological replicates. The proteins indicated in red had a second-generation p value of 0. The vertical line indicates a log_2_ fold change (log_2_FC) of 0. (**B**) Pathway analysis indicating the top suppressed pathways in UGCG-deficient CD8^+^ T cells from Figure 6A. NES, normalized enrichment score. (**C-D**) Quantification of membrane lipid rafts as determined by cholera toxin B (CTxB) binding. Representative histogram (*left*) and quantification of relative geometric mean fluorescence intensity (MFI; *right*) of CTxB expression on (C) shCtrl- and sh*Ugp2*-expressing or (D) NTC- and sg*Ugcg*-modified CD8^+^ T cells. Data represent the mean ± SEM (n = 3-4). (**E**) Lipid raft aggregation following TCR crosslinking in NTC- and sg*Ugcg*-modified CD8^+^ T cells. *Left*, Representative confocal images of an NTC- and sg*Ugcg*-modified cell stained with CTxB (green), streptavidin (magenta), and DAPI (blue) after incubation with anti-CD3ε antibody and crosslinking with biotin-labeled IgG. *Right*, Quantification of lipid raft (CTxB) aggregation and CD3ε intensity (normalized to cell size) in NTC- (n = 46) and *sgUgcg*-modified (n = 12) cells was conducted 30 min after crosslinking. Data represent the mean ± SD. Statistical significance was determined using Mann-Whitney tests. AU, arbitrary unit.

GSLs, particularly gangliosides, are integral components of lipid rafts, which are lipid-rich microdomains at the plasma membrane that serve as platforms for cell signaling^36^. In T cells, stimulation of the TCR is followed by aggregation of lipid rafts, which brings together key proteins involved in TCR signal transduction^37^. Importantly, lipid raft defects have been associated with compromised T cell proliferation^38^. Hence, we explored lipid raft aggregation as a putative mechanism for the reduced cell proliferation triggered by loss of glucose-dependent GSL biosynthesis (**Figure 5**). To determine the impact of *Ugp2* silencing or *Ugcg* deletion on lipid rafts in CD8^+^ T cells, we took advantage of the cholera toxin B subunit (CTxB), which binds GM1 and other gangliosides and is used to measure lipid rafts at the cell surface^39,40^. Indeed, both UGP2-depleted and UGCG-deficient CD8^+^ T cells displayed a reduction in the MFI of CTxB, indicating reduced levels of lipid rafts at the plasma membrane (**Figures 6C-D**). Moreover, while TCR crosslinking resulted in lipid raft aggregation in control cells, UGCG-deficient CD8^+^ T cells displayed impaired lipid raft aggregation (**Figure 6E**). Together, these data indicate that glucose-dependent GSL biosynthesis is required to maintain the ganglioside composition of lipid rafts in T cells.

### UGCG controls CD8^+^ T cell cytotoxic function and anti-tumor immunity

Given the decrease in cytotoxic factors and lipid raft defects in UGCG-deficient CD8^+^ T cells (**Figure 6A-B**), we evaluated the role of glucose-dependent GSL biosynthesis in T cell-mediated tumor control. First, we evaluated the cytolytic capacity of UGCG-deficient CD8^+^ T cells. Control or UGCG-deficient Thy1.1^+^CD8^+^ OT-I cells were transferred into Thy1.2^+^ C57BL/6J hosts, infected with *Lm*-OVA, and the cytotoxic activity of antigen-specific OT-I cells isolated from infected animals at 7 dpi was assessed via T cell killing assay using OVA-expressing MC38 (MC38-OVA) colon cancer cells as targets^16^. UGCG-deficient OT-I cells were impaired in their ability to lyse cancer cells, with ∼6x the number of UGCG-deficient cells required to kill the same number of cancer cells as control OT-I cells (**Figure 7A**). Moreover, at a fixed effector:target ratio, UGCG-deficient OT-I cells displayed a reduced rate of T cell killing (**Figure 7B**). Consistent with our proteomics analysis (**Figure 6A-B**) and the reduced killing capacity of UGCG-deficient CD8^+^ T cells (**Figure 7A-B**), physiologically activated UGCG-deficient CD8^+^ OT-I cells had lower Granzyme B (GZMB) expression *ex vivo* (**Figure 7C**). Thus, in addition to proliferative defects, CD8^+^ T cells unable to maintain GSL production from glucose display impaired cytotoxic function.

**Figure 7:**
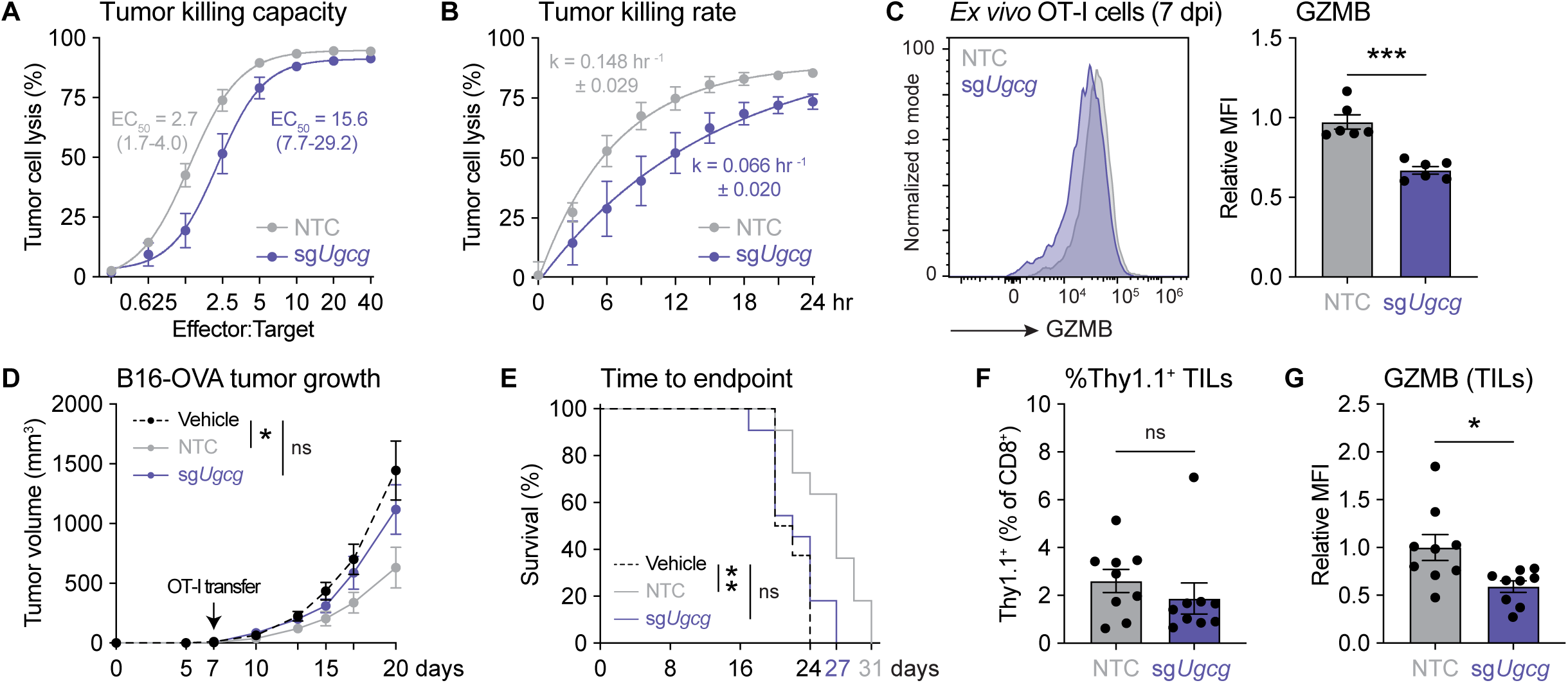
UGCG controls CD8^+^ T cell cytotoxic function and anti-tumor immunity. (**A-B**) Cytolytic activity of UGCG-deficient CD8^+^ T cells. (A) Tumor cell kill curve for control (NTC) and UGCG-deficient (sg*Ugcg*) CD8^+^ OT-I cells isolated from *Lm*-OVA-infected mice at 7 days post-infection (dpi). OT-I cells were co-cultured *ex vivo* with MC38-OVA cancer cells at the indicated effector:target (E:T) ratio and cancer cell viability measured after 24 h. The E:T ratio required to kill 50% of cancer cells (EC_50_) and 95% confidence interval are indicated for each genotype. Data represent the mean ± SEM (n = 6). (**B**) Timecourse of cancer cell lysis for the experiment in (A) at a fixed effector:target ratio of 10:1. The rate constant (k) ± SEM is indicated for each genotype. Data represent the mean ± SEM (n = 6). (**C**) Granzyme B (GZMB) production in NTC- and sg*Ugcg*-modified CD8^+^ OT-I cells. OT-I cells isolated from *Lm*-OVA-infected mice at 7 dpi as in (A) were stained for intracellular GZMB expression. Representative histogram (*left*) and quantification of relative mean fluorescence intensity (MFI; *right*) of GZMB in NTC- and sg*Ugcg*-modified CD8^+^ T cells. Data represent the mean ± SEM (n = 6). (**D**) Timecourse of B16-OVA melanoma tumor growth in mice that received NTC- or sg*Ugcg*-modified CD8^+^ OT-I cells. OT-I cells (1×10^6^) were transferred into tumor-bearing mice 7 days post-implantation (n = 11 mice/group). Vehicle control-injected mice (n = 8) were included as a “no adoptive transfer” control. Statistical significance was determined using a mixed-effects model with Geisser-Greenhouse correction and Dunnett’s multiple comparisons test. (**E**) Kaplan-Meier curve of time-to-humane endpoint for tumor-bearing mice from (D). Statistical significance was determined using log-rank tests and a Bonferroni correction for multiple comparisons. (**F**) Percentage of antigen-specific (Thy1.1^+^) NTC- and sg*Ugcg*-modified CD8^+^ OT-I cells in B16-OVA tumors 7 days post-adoptive transfer. Data represent the mean ± SEM (n = 9 mice/group). (**G**) GZMB production in B16-OVA TILs from (F). Relative MFI of GZMB in Thy1.1^+^CD8^+^ NTC- and sg*Ugcg*-modified OT-I cells 7 days post-adoptive transfer. Data represent the mean ± SEM (n = 9 mice/group).

Finally, to test whether cytotoxic defects linked to impaired GSL biosynthesis impact anti-tumor immunity, we adoptively transferred control or UGCG-deficient Thy1.1^+^CD8^+^ OT-I cells into B16-OVA tumor-bearing mice and measured tumor growth over time. While transfer of control OT-I cells significantly delayed tumor growth and prolonged overall survival, animals that received UGCG-deficient OT-I cells displayed similar tumor growth as mice that received no OT-I cells (**Figure 7D-E**). Unlike the *Lm*-OVA infection model (**Figure 5I-J**), the defect in tumor control conferred by UGCG-deficient CD8^+^ T cells was not due to a reduction in the percentage of antigen-specific CD8^+^ T cells in the tumor (**Figure 7F**). Rather, we observed a significant reduction in GZMB expression in antigen-specific, UGCG-deficient tumor-infiltrating lymphocytes (TILs) (**Figure 7G**). Collectively, our data indicate that glucose-dependent GSL biosynthesis is required for optimal CD8^+^ T cell cytotoxic function and effective tumor control.

## DISCUSSION

Glucose is an essential substrate for T cell proliferation and function. While glucose is a prominent bioenergetic substrate for ATP production in most cell types, it has become increasingly clear that energy production is not the only metabolic fate for glucose important for T cell responses *in vivo*. Here, we show that nucleotide sugar biosynthesis—specifically the production of UDP-Glc—is an essential fate for glucose in proliferating CD8^+^ Teff cells independent of energy production. The rapid production of UDP-Glc from glucose is a feature of both mouse and human CD8^+^ T cells and requires the coordination of multiple glucose-fueled anabolic pathways (i.e., PPP, nucleotide biosynthesis, aspartate biosynthesis) for optimal production. Glucose-dependent UDP-Glc biosynthesis is critical for both quiescent T cell homeostasis and Teff cell expansion. Conditional deletion of *Ugp2* in T cells does not notably affect thymic development, but significantly depletes mature T cell populations in the periphery. Concordantly, preventing the conversion of glucose to UDP-Glc by depleting UGP2 in activated CD8^+^ T cells cripples their ability to respond to *Lm*-OVA infection *in vivo*. Importantly, blocking UDP-Glc biosynthesis did not affect the contribution of glucose to serine or nucleotide biosynthesis, which we previously showed are important for T cell expansion^9,41^, nor did it impair TCA cycle metabolism. While UDP-Glc is a substrate for several metabolic pathways, we found that blocking GSL biosynthesis (via UGCG inhibition) was sufficient to phenocopy the defects we observed with UGP2 depletion, implicating GSL biosynthesis as the key metabolic fate of glucose critical for optimal CD8^+^ Teff cell responses. Mechanistically, glucose-dependent GSL biosynthesis is essential for maintaining membrane lipid raft integrity, and disrupting this pathway led to reduced CD8^+^ T cell cytotoxicity and poor tumor control. Taken together, our results establish glucose-dependent GSL biosynthesis as an essential metabolic pathway in CD8^+^ T cells that is independent of other known biosynthetic and bioenergetic fates of glucose.

Contrary to current models, our findings suggest that glucose functions predominantly as a biosynthetic precursor, rather than a bioenergetic substrate, in CD8^+^ T cells under physiologic conditions. This is supported by recent evidence demonstrating that, under physiologic conditions, increased TCA cycle metabolism and oxidative ATP production by activated T cells is fueled by other, non-glucose carbon sources, including glutamine^15,17^, lactate^15^, acetate^17,42,43^, and ketone bodies^15,16,44^. In the absence of these non-glucose carbon sources (i.e., *in vitro* cell culture), excess glucose can be used by proliferating CD8^+^ T cells as both a biosynthetic and bioenergetic substrate. However, under physiologic conditions where these other carbon sources are abundant^45–47^, glucose is spared from entering the TCA cycle and instead can feed biosynthetic pathways that generate other metabolites needed for cell proliferation, including serine^9,41^, nucleotides^9,17,41^, and UDP-GlcNAc^48^. Notably, these glucose-dependent metabolic pathways all converge on the synthesis of GSLs, which we show here are essential for CD8^+^ Teff cell expansion and cytotoxic function. After only 2 h of ^13^C-glucose infusion in *Listeria*-infected mice, almost 20% of the HexCer pools in activated CD8^+^ Teff cells were labeled from ^13^C-glucose, indicating rapid turnover of these lipid pools *in vivo*. Depleting GSL pools in activated T cells (via UGCG deletion) led to a loss of lipid raft integrity at the plasma membrane. Thus, sparing glucose from being metabolized for ATP production allows proliferating T cells to direct glucose towards GSL synthesis *in vivo* and maintain these essential lipid pools. We posit that the metabolic processes that fuel *in vivo* CD8^+^ Teff cell activation, proliferation, and function are partitioned, whereby glutamine and other physiologic carbon sources such as ketone bodies are preferentially oxidized via the TCA cycle to fuel T cell bioenergetics, which spares glucose to fuel biosynthetic pathways that branch from glycolysis, such as nucleotide, glycan, and GSL biosynthesis. How glucose-dependent GSL biosynthesis changes over the course of an immune response and whether GSLs contribute to T cell homeostasis during quiescent cell states remains to be determined.

Glycobiology remains an important but poorly understood area of immunology, due in part to the complex structures and biosynthesis of glycans, and the wide range of biological processes they regulate. T cell activation is associated with dramatic glycocalyx remodeling^49,50^, which has implications for both T cell activation and function^51–53^. Our data reveal that a major fate of glucose in both mouse and human activated CD8^+^ T cells is the rapid biosynthesis of nucleotide sugars required for glycosylation reactions, including UDP-GlcNAc and UDP-Glc. We show that inhibiting UDP-Glc biosynthesis via UGP2 depletion blunts N-glycan extension and impairs GSL biosynthesis, implicating UGP2 as a central regulator of the glycocalyx in CD8^+^ T cells. In the present study, we focused primarily on the role of UDP-Glc in GSL biosynthesis. We demonstrated, for the first time, that CD8^+^ Teff cells actively synthesize GSLs from glucose *in vivo* in response to pathogen challenge. In particular, we observed increased pools and ^13^C-glucose labeling of asialo-series (i.e., GA1) gangliosides in proliferating Teff cells compared to resting Tn cells. This observation is consistent with a previous study that found mouse CD8^+^ T cells predominantly express asialo-series gangliosides, whereas mouse CD4^+^ T cells express a-series gangliosides (e.g., GM3-derived GM2)^54^. Interestingly, in human CD8^+^ T cells, we detected glucose-derived GM3 rather than GA1 gangliosides, suggesting that while both mouse and human activated CD8^+^ T cells synthesize GSLs, the types of GSLs produced may be species-dependent. Concordantly, a recent comprehensive indexing of glycan structures in activated mouse and human T cells revealed that increased glycan biosynthesis is a common feature of T cell activation, but the specific glycan structures expressed in mouse and human T cells differed due to divergent expression of multiple glycosyltransferase enzymes^55^. Species-dependent differences in glycan synthesis may explain key differences in immune responses observed across species.

A key consequence of inhibiting GSL biosynthesis in CD8^+^ T cells (via UGP2 depletion or UGCG deletion) was a reduction in ganglioside-rich lipid rafts at the plasma membrane. We further showed that lipid raft aggregation following TCR crosslinking is compromised in UGCG-deficient CD8^+^ T cells. Lipids rafts have been implicated in T cell proliferation^54^, polarization and migration^56^, responsiveness to cytokines^57^, and formation of the cytotoxic immunological synapse^58^. We therefore propose a model whereby glucose-dependent GSL biosynthesis tethers nutrient availability to the maintenance of signaling complexes at the plasma membrane, ensuring T cell proliferation and cytotoxic function are supported by sufficient glucose levels. This model is supported by our proteomics data, which showed reduced expression of granzymes and proteins involved in actin cytoskeleton organization in UGCG-deficient CD8^+^ T cells. Moreover, lipid rafts have been reported to control naïve CD8^+^ T cell homeostasis through tonic TCR signaling^57^, and therefore a defect in lipid rafts could explain the reduced number of peripheral T cells we observed in *Ugp2*^fl/fl^*Cd4*-*Cre* mice. Data from multiple different cell types, including the data we present here, show that inhibiting GSL biosynthesis *in vitro* impairs cell proliferation, but has little effect on cell viability^59,60^; however, *in vivo*, GSLs are key mediators of cell-cell and cell-environment communication, which are important for coordinating complex biological processes including embryonic development^60^ and immune responses^54^. In addition to their structural contribution to lipid rafts, membrane GSLs may also regulate cell signaling and cell-cell adhesion via carbohydrate-carbohydrate interactions^55^.

Glucose-dependent GSL biosynthesis has emerged as a key metabolic feature of tumor cells. For example, in oncogenic KRAS-driven pancreatic cancer cells, glucose-dependent GSL biosynthesis regulates KRAS plasma membrane localization and signaling^61^. Other studies show that cancer cell-intrinsic GSL biosynthesis promotes tumor growth by mediating immune evasion, and that inhibiting GSL biosynthesis in cancer cells increases cancer cell sensitivity to IFN-γ-induced growth arrest^62^ and improves antigen presentation to cytotoxic CD8^+^ T cells^63^. In this vein, eliglustat treatment may synergize with immune checkpoint blockade therapy to inhibit tumor growth^62,63^. Here, our findings suggest that systemic treatments that target UGCG, such as eliglustat, may compromise the cytotoxic capacity of CD8^+^ T cells and hinder their ability to control tumor growth. Hence, while short-term eliglustat treatment may reduce tumor burden by inhibiting cancer cell-intrinsic GSL biosynthesis, prolonged treatment may have unintended consequences for long-term anti-tumor immunity. By contrast, immunometabolic modulation of GSL biosynthesis may prove effective in modulating pathology in diseases driven by pathogenic T cells, such as autoimmune diseases.

## LIMITATIONS OF THE STUDY

Given that deletion of *Ugp2* during T cell development was detrimental to T cell homeostasis and that surviving peripheral T cells still expressed UGP2, we were unable to study how disrupting glucose-dependent UDP-Glc and GSL biosynthesis affects the transition from the Tn to Teff cell state. We instead focused on studying how these metabolic pathways impact proliferating Teff cells *in vivo*. Inducible *Ugp2* or *Ugcg* deletion models will enable the study of GSL metabolism during different T cell state transitions in the future. Our data establish that glucose is used by CD8^+^ T cells to sustain GSL biosynthesis during an immune response; however, further research is required to determine how different GSLs expressed at the plasma membrane of T cells interact with other membrane components on the same cell or neighboring cells to coordinate T cell responses.

## ACKNOWLEDGMENTS

We thank Dr. Jim Dennis and members of the Jones and Krawczyk laboratories for scientific discussions contributing to this manuscript. We thank Jade Desjardins, Mitra Cowan, and the McGill Integrated Core for Animal Modeling for generating the *Ugp2*-floxed mouse model. We thank Jeanie Wedberg and Margene Brewer for administrative assistance. We thank members of the Van Andel Institute (VAI) Core Facilities for technical assistance, including Mass Spectrometry (RRID: SCR_024903), Flow Cytometry (RRID: SCR_022685), Bioinformatics and Biostatistics (RRID: SCR_024762), Vivarium (RRID: SCR_023211), Pathology and Biorepository (RRID: SCR_022912) and Optical Imaging (RRID: SCR_021968). We especially thank Christine Isaguirre and Molly Hopper (Mass Spectrometry), Rachael Sheridan (Flow Cytometry), Lisa Turner (Pathology and Biorepository), and Lorna Cohen (Optical Imaging) for their technical support. J.L. is supported by a VAI Metabolism & Nutrition (MeNu) Program Pathway-to-Independence Award and Canadian Institutes of Health Research (CIHR) Fellowship (MFE-181903). M.S.D. is supported by a VAI MeNu Program Pathway-to-Independence Award, Fonds de recherche du Québec-Santé (FRQS) Postdoctoral Fellowship (0000289124), and CIHR Fellowship (MFE-403514). C.M.K. is supported by the National Institute of Allergy and Infectious Diseases (NIAID, R21AI153997) and VAI. B.B.H. is supported by VAI. R.G.J. is supported by the Paul G. Allen Frontiers Group Distinguished Investigator Program, Chan Zuckerberg Initiative (CZI), NIAID (R01AI165722), and VAI.

## AUTHOR CONTRIBUTIONS

Conceptualization, J.L. and R.G.J.; Experimental Design, J.L., B.M.O., A.R-O., K.L.G., C.R.E., C.M.K., R.D.S., and R.G.J.; Investigation, J.L., L.M.D., B.M.O., R.T., A.R-O., M.S.D., A.E.E., S.E.C., and H.L.; Resources, D.G.R.; Data Analysis, J.L., R.T., A.R-O., A.E.E., M.P.V., H.D., K.L.G., C.D.C., Z.B.M., H.L., R.D.S., and R.G.J.; Writing, J.L. and R.G.J.; Visualization, J.L. and K.S.W.; Supervision, C.R.E., B.B.H., R.D.S., and R.G.J.; Funding Acquisition, R.G.J.

## DECLARATION OF INTERESTS

R.G.J. is a scientific advisor to Servier Pharmaceuticals and is a member of the Scientific Advisory Board of Immunomet Therapeutics.

## STAR METHODS

### KEY RESOURCES TABLE

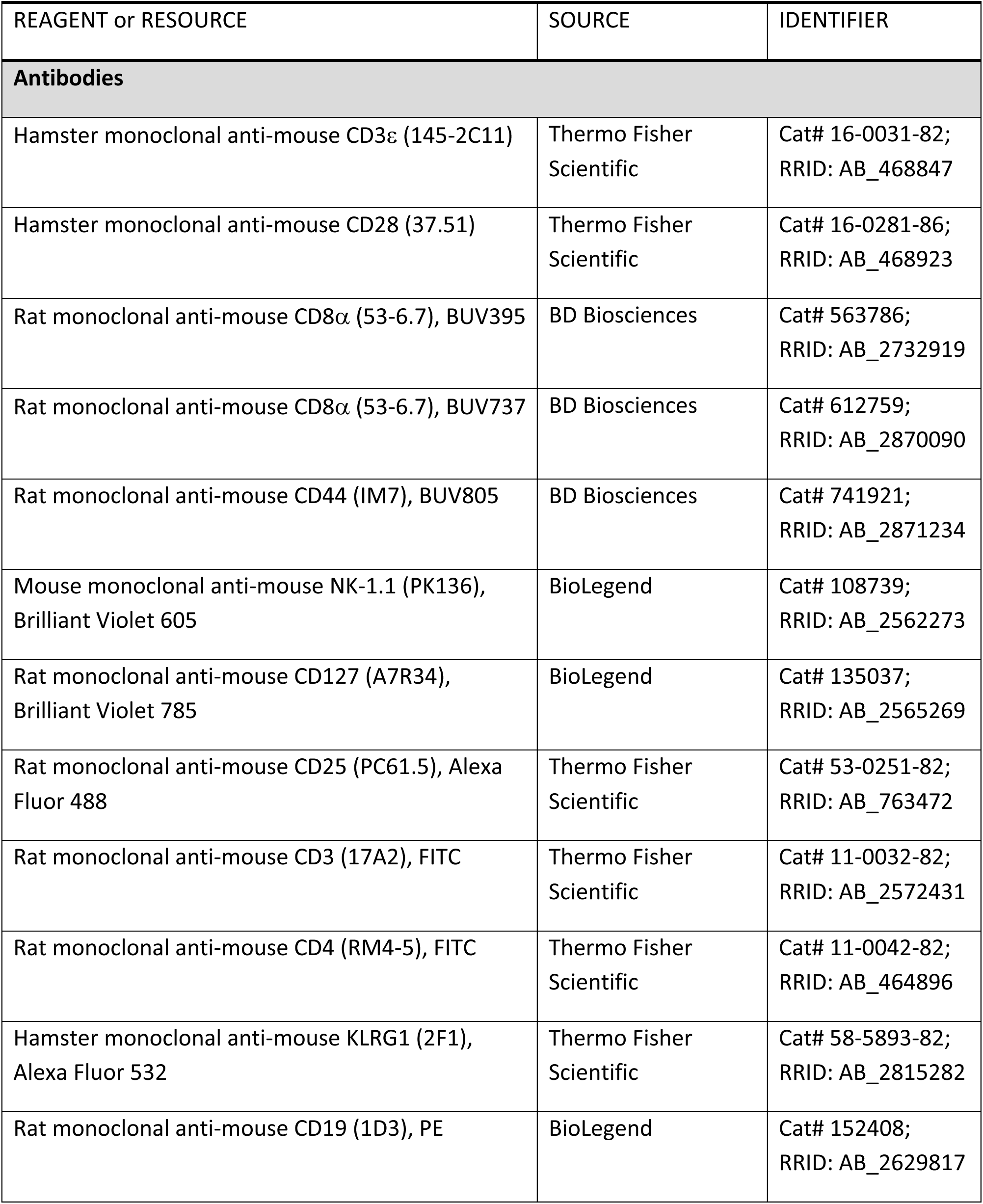
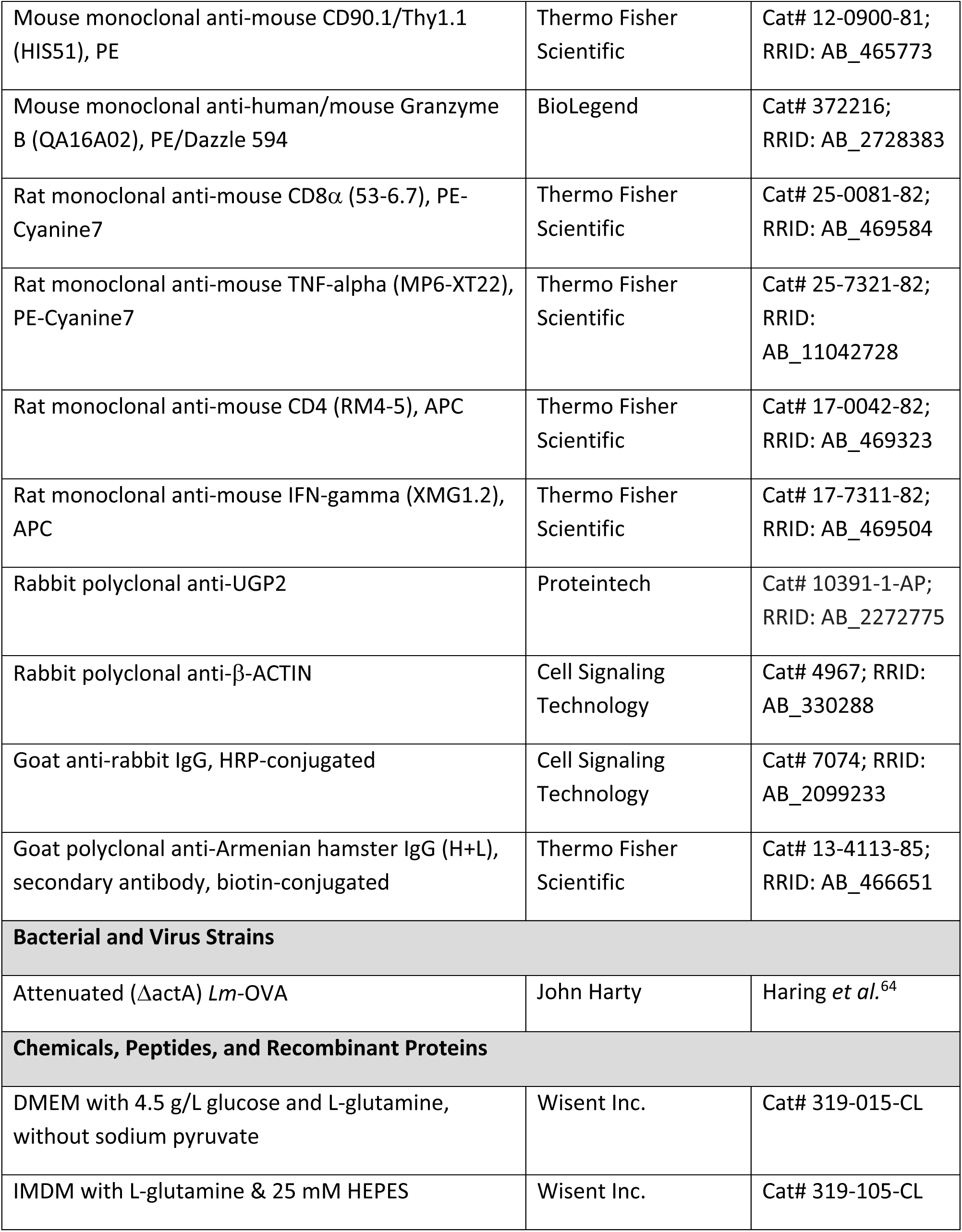
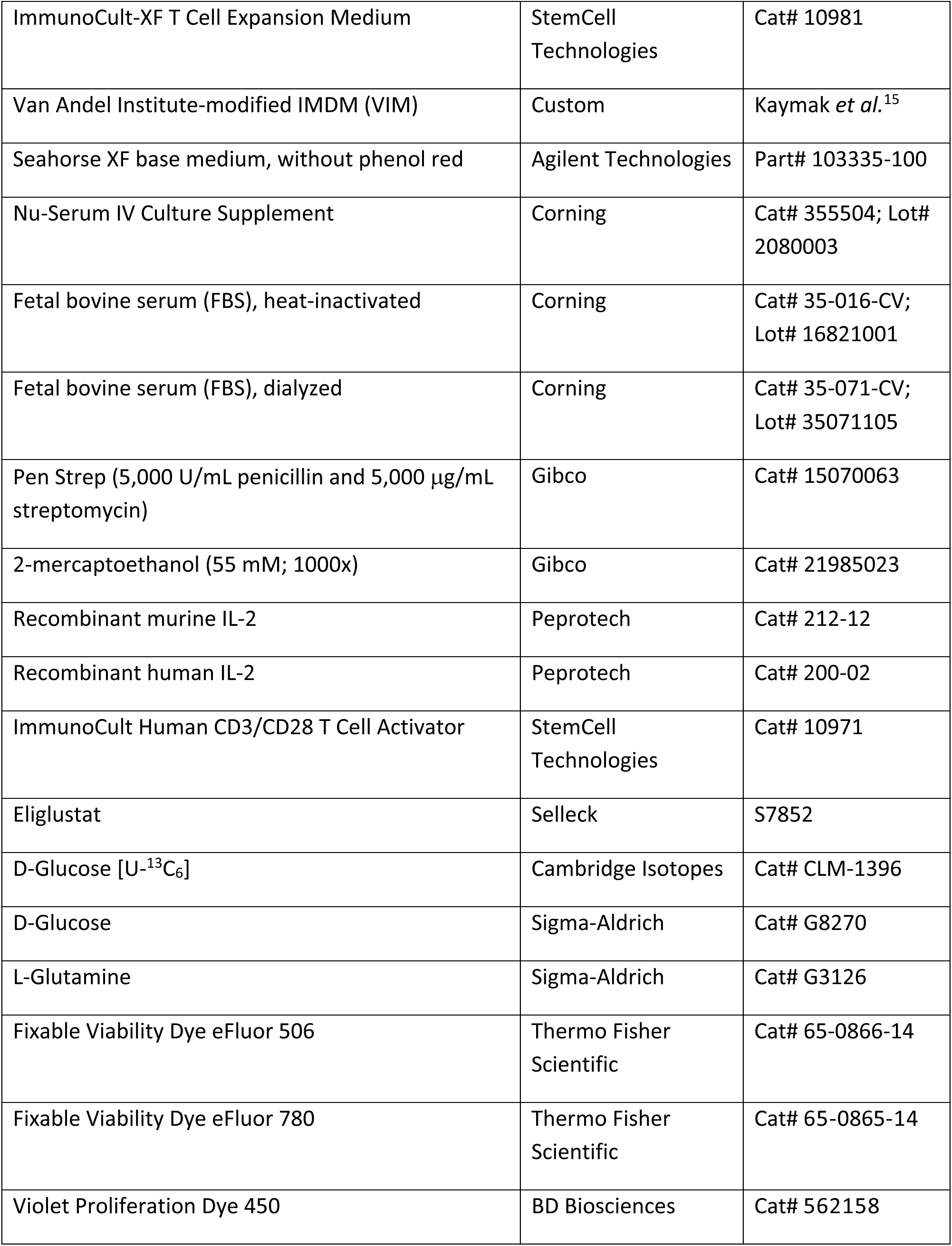
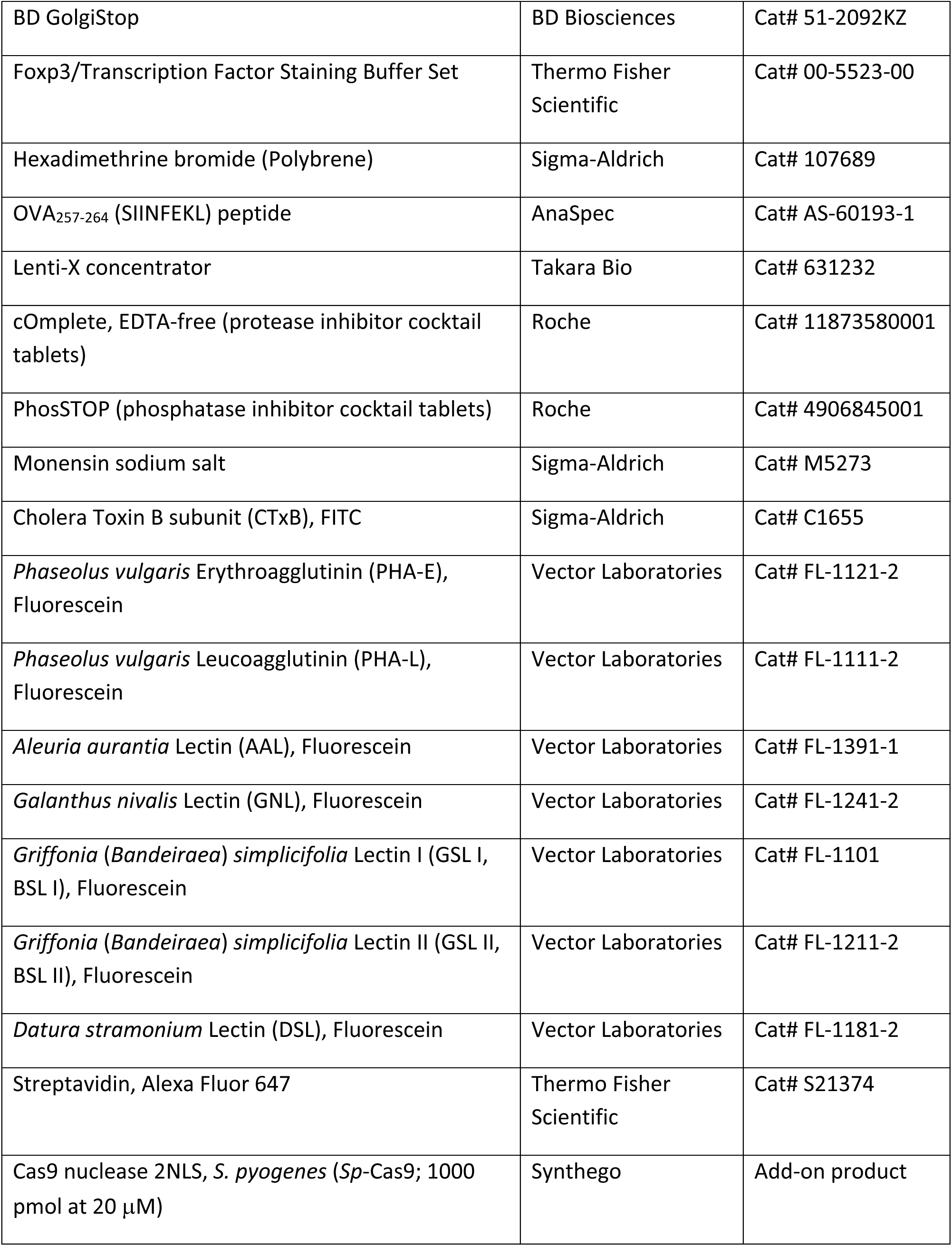
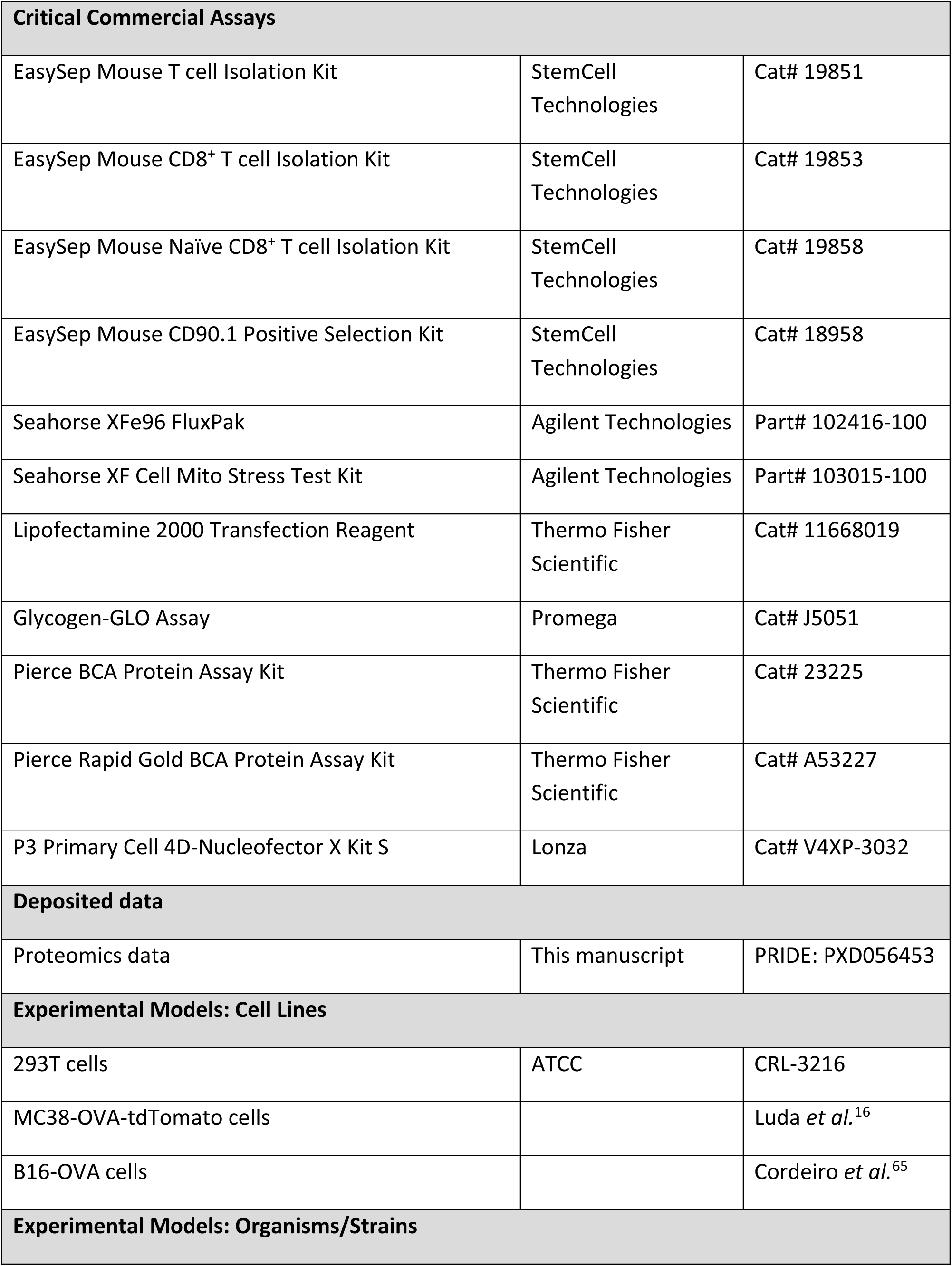
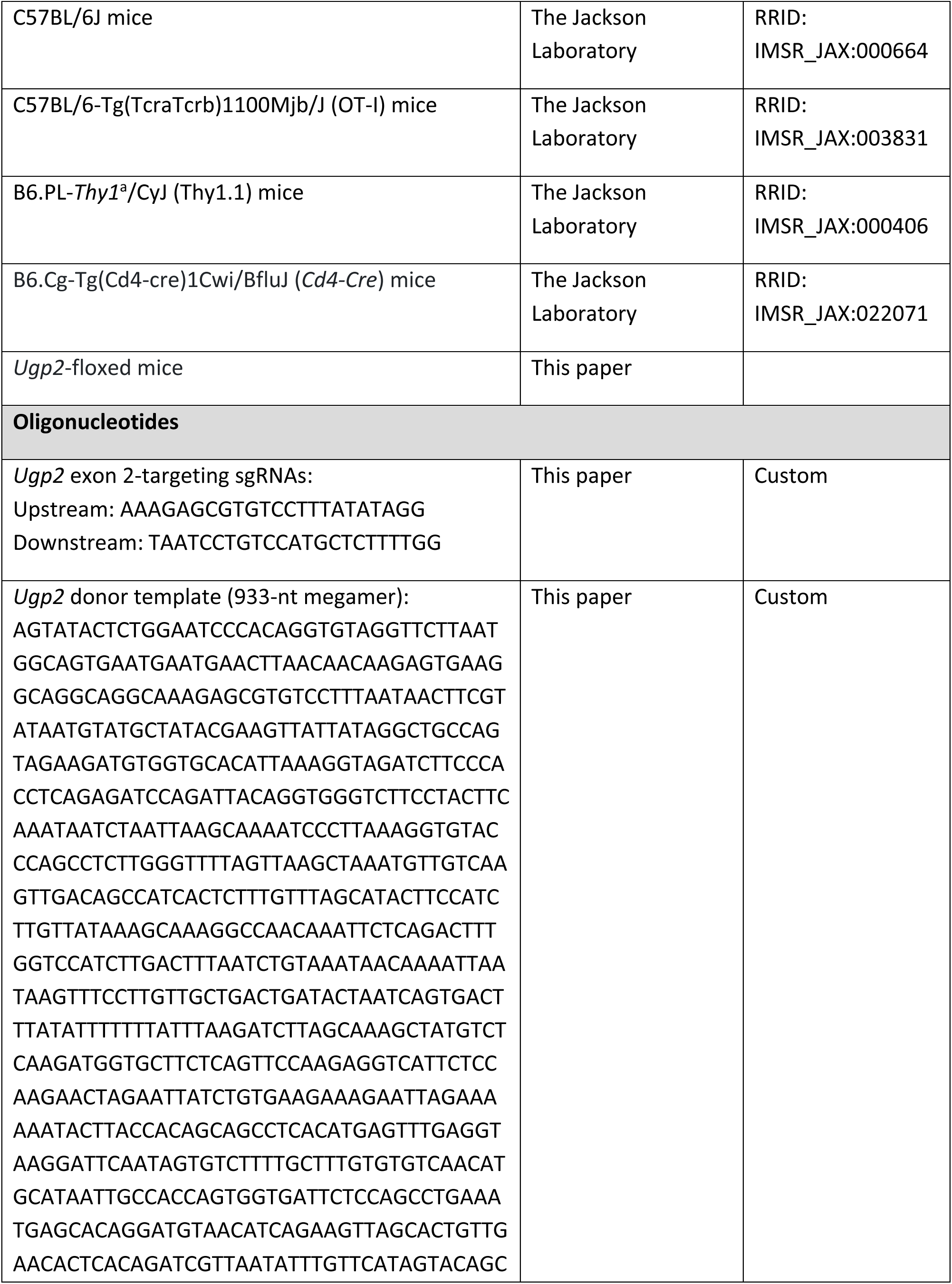
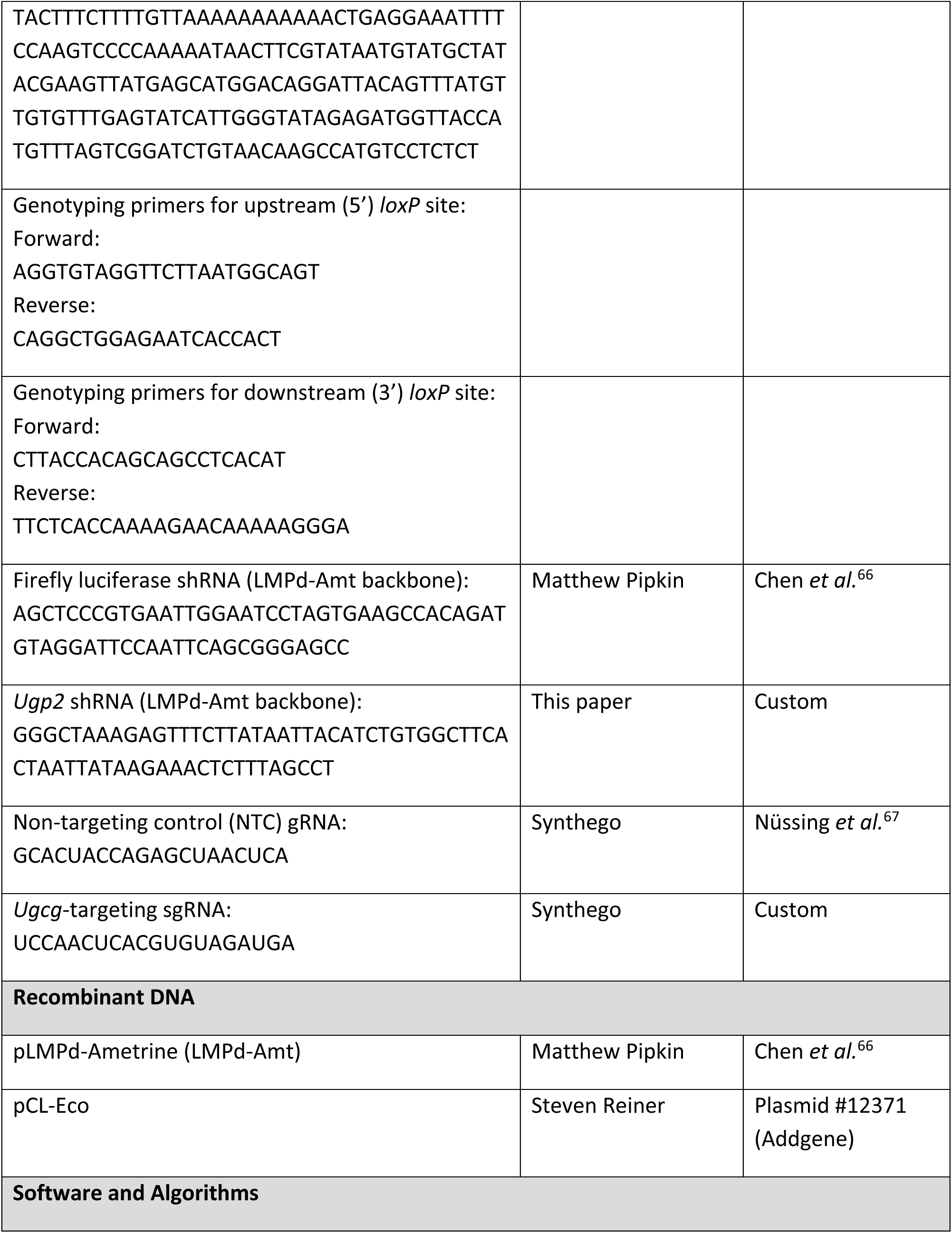
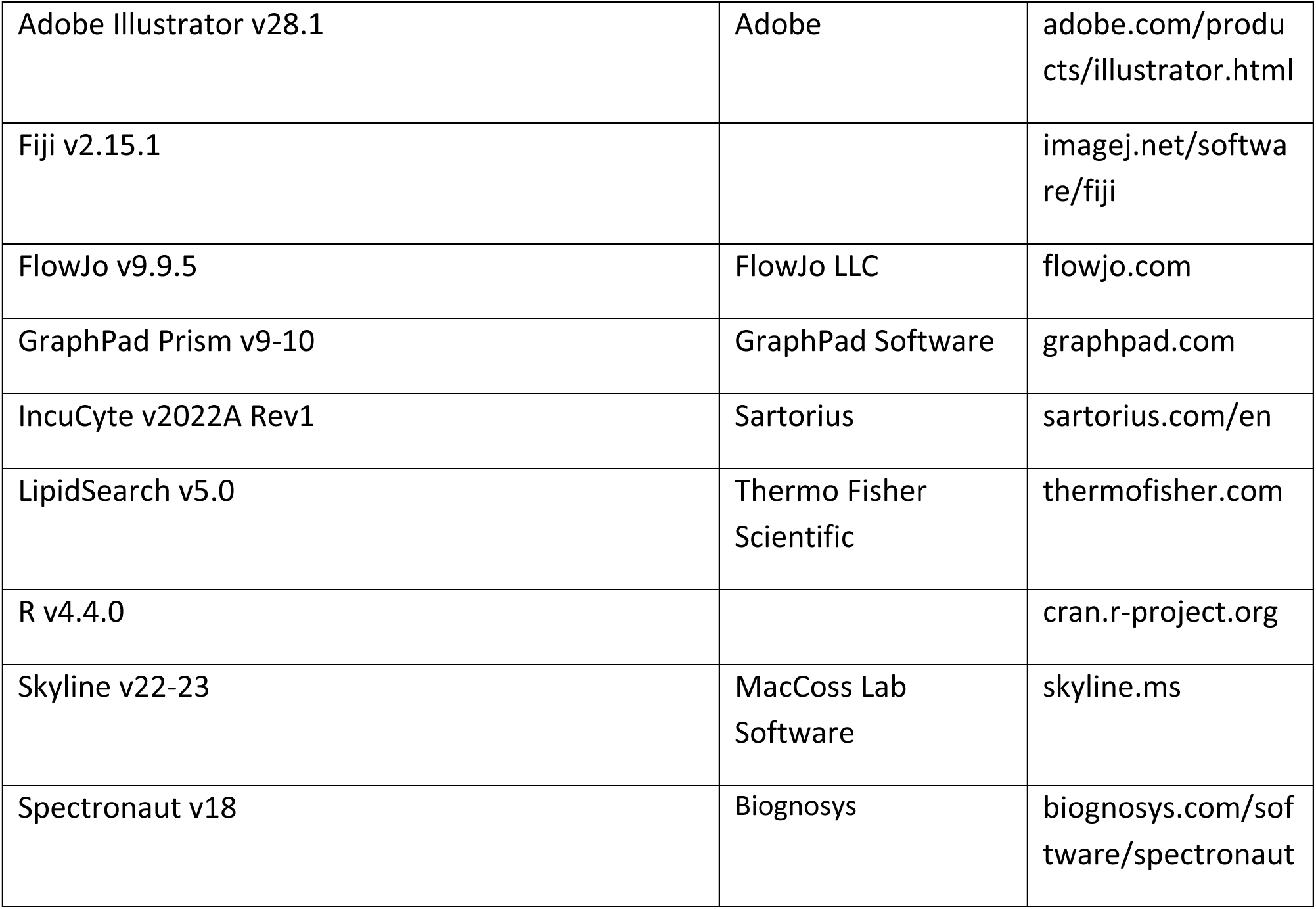

### RESOURCE AVAILABILITY

#### Lead Contact

Additional information and requests for raw data or reagents should be directed to, and will be made available by, the corresponding author, Dr. Russell G. Jones.

#### Materials availability

All unique, stable reagents generated in this study will be made available from the Lead Contact with a completed Material Transfer Agreement. The LMPd-Amt-sh*Ugp2* plasmid will be deposited to Addgene.

#### Data and code availability

The mass spectrometry proteomics data have been deposited to the ProteomeXchange Consortium via the PRIDE partner repository with the dataset identifier PXD056453. The proteomics data and confocal image analysis pipelines are hosted on GitHub (see METHOD DETAILS for links).

### EXPERIMENTAL MODEL AND SUBJECT DETAILS

#### Mice

C57BL/6J (RRID: IMSR_JAX:000664), B6.PL-*Thy1*^a^/CyJ (Thy1.1; RRID: IMSR_JAX:000406), C57BL/6-Tg(TcraTcrb)1100Mjb/J (OT-I; RRID: IMSR_JAX:003831), and B6.Cg-Tg(Cd4-cre)1Cwi/BfluJ (*Cd4*-*Cre*; RRID: IMSR_JAX:022071) mice were purchased from The Jackson Laboratory. Homozygous Thy1.1 mice were crossed to homozygous OT-I mice to generate heterozygous Thy1.1 OT-I mice, which were used to isolate Thy1.1^+^CD8^+^ OT-I cells. All mice were bred and maintained in grouped housing at the Van Andel Institute (VAI) Vivarium under specific pathogen-free conditions. All procedures involving mice were completed as recommended in the Guide for the Care and Use of Laboratory Animals, and all protocols were approved by VAI’s Institutional Animal Care and Use Committee (IACUC). Genotyping was performed using DNA extracted from ear biopsies and defined primer sets^16^ (see **Key Resource Table**). Experiments were performed using female mice between 8-12 weeks of age, unless otherwise indicated.

#### Cell lines

293T cells (CRL-3216), MC38 murine colon cancer cells expressing OVA and tdTomato (MC38-OVA-tdTomato)^16^, and B16-F10 murine melanoma cells expressing OVA (B16-OVA)^65^ were cultured in Dulbecco’s Modified Eagle’s Medium (DMEM) (Wisent Inc.) supplemented with 10% heat-inactivated fetal bovine serum (FBS), 1% penicillin-streptomycin (Gibco; [final] = 50 U/mL penicillin and 50 μg/mL streptomycin), and L-glutamine at a final concentration of 6 mM. 293T and MC38-OVA-tdTomato cells are female, and B16-OVA cells are male. All cells were cultured at 37°C in a humidified 5% CO_2_ incubator.

#### Primary cells

Frozen human peripheral blood CD8^+^ T cells from three different healthy donors were purchased from StemCell Technologies (Cat# 200-0164). Two donors were female (Lot# 2210425008 and 2210427003) and one donor was male (Lot# 2210403009). Cells were cultured in ImmunoCult-XF T Cell expansion Medium (StemCell Technologies) supplemented with 50 U/mL penicillin and 50 μg/mL streptomycin (Gibco) and 300 U/mL recombinant human IL-2 (Peprotech) at 37°C in a humidified 5% CO_2_ incubator. Cells were activated for 72 h with ImmunoCult Human CD3/CD28 T Cell Activator (StemCell Technologies; 25 μL per 1×10^6^ cells in 1 mL final volume).

### METHOD DETAILS

#### Generation of Ugp2-floxed mice

*Ugp2*-floxed animals were generated by CRISPR/Cas9 gene editing at the McGill Integrated Core for Animal Modeling, as previously described^68^. Single gRNA (sgRNA) sequences flanking *Ugp2* exon 2 were purchased from Synthego and the single-stranded megamer (933-nt) donor template was purchased from Integrated DNA Technologies (IDT; see **Key Resources Table**). The sgRNAs (25 ng/μL each) were complexed with Alt-R *Sp*-Cas9 Nuclease V3 (IDT, Cat# 1081059; 50 ng/μL) and microinjected simultaneously with the donor template (20 ng/μL) into the pronucleus of zygotes from C57BL/6J mice. After microinjection, the embryos were implanted into pseudo-pregnant CD-1 female mice for gestation, as described previously^68^. Successful *loxP* insertion was confirmed using PCR primers flanking the mutation sites (see **Key Resources Table**) followed by Sanger sequencing. *Ugp2*-floxed mice were crossed to hemizygous *Cd4*-*Cre* mice to generate *Ugp2*^fl/fl^*Cd4*-*Cre* mice. Cre-negative *Ugp2*^fl/fl^ mice were used as controls.

#### T cell purification and culture

For mouse T cell isolations, total (CD3^+^) T cells, total CD8^+^ T cells, or naïve CD8^+^ T cells were purified from the spleen and peripheral lymph nodes by negative selection (StemCell Technologies). Cells were cultured in Iscove’s Modified Dulbecco’s Medium (IMDM; Wisent Inc.) supplemented with 10% Nu-Serum IV culture supplement (Corning), 50 U/mL penicillin and 50 μg/mL streptomycin (Gibco), 50 μM 2-mercaptoethanol (Gibco), and 2 mL L-glutamine (Gibco; 6 mM final concentration). T cells (1×10^6^ cells/mL) were activated *in vitro* via stimulation with plate-bound anti-CD3ε (clone 145-2C11; 2 μg/mL) and anti-CD28 (clone 37.51; 1 μg/mL) antibodies for 48-72 hr. After activation, cells were propagated in IMDM supplemented with 50 U/mL recombinant murine IL-2 (Peprotech). Proliferating cells were re-seeded at 4×10^5^ cells/mL in fresh medium and IL-2 every two days.

#### shRNA cloning, retrovirus production, and viral transduction

Short hairpin RNAs (shRNAs) against firefly luciferase (shCtrl) or *Ugp2* (sh*Ugp2*; see **Key Resources Table** for sequences) were subcloned into the pLMPd-Amt vector between the XhoI and EcoRI restriction enzyme sites^66^. 293T cell transfection, retrovirus production, and viral transduction of activated CD8^+^ T cells were performed as previously described^16^.

#### CRISPR/Cas9 gene editing in primary CD8^+^ T cells

A singe guide RNA (sgRNA) targeting the murine *Ugcg* gene (5’–UCCAACUCACGUGUAGAUGA–3’) and a non-targeting control (NTC) sgRNA (5’–GCACUACCAGAGCUAACUCA–3’)^67^ were obtained from Synthego (CRISPRevolution sgRNA EZ Kit). For sgRNA/Cas9 ribonucleoprotein (RNP) complex formation, 1.86 μL of *Sp*-Cas9 nuclease (Synthego) was combined with 3.4 μL of target sgRNA (0.1 nmol/μL stock in nuclease-free water) and incubated for 10 min at room temperature. In the meantime, 2.5×10^6^ activated CD8^+^ T cells were pelleted, washed twice with phosphate-buffered saline (PBS), and resuspended in 20 μL of P3 solution mix (P3 Primary Cell 4D-Nucleofector X Kit S, Lonza). After RNP complex formation, the cells were added to the RNP complex mixture, and the RNP/cell mixture was then transferred to a well of a 16-well Nucleocuvette Strip (Lonza). Cells were electroporated using a 4D-Nucleofector (Lonza; program DN 100). Immediately after electroporation, 150 μL of 37°C pre-warmed IMDM (antibiotic-free) was added to the cells, which were then left to rest at 37°C for 10 min. Cells were transferred to 2.5 mL of IMDM (antibiotic-free) in a well of a 12-well plate. The cells were used for adoptive transfer or other experiments 2-3 days post-electroporation. Knockout of *Ugcg* was confirmed by LC-MS and/or CTxB staining.

#### Adoptive transfer and infection with L. monocytogenes (Lm-OVA)

Mice were immunized intravenously with a sublethal dose of recombinant attenuated *Listeria monocytogenes* expressing OVA (*Lm*-OVA; 2×10^6^ colony-forming units (CFU)), as previously described^16,69^. For adoptive transfer experiments involving immunophenotyping at 7 days post-infection (dpi), 5×10^3^ Thy1.1^+^CD8^+^ OT-I cells were injected intravenously into Thy1.2^+^ C57BL/6J mice, followed by *Lm*-OVA infection 1 day later. Splenocytes were isolated from mice at 7 dpi and analyzed *ex vivo* by flow cytometry. For *in vivo* ^13^C-glucose infusion or *ex vivo* ^13^C-glucose tracing experiments, Thy1.2^+^ C57BL/6J mice received 2×10^6^ Thy1.1^+^CD8^+^ OT-I cells intravenously and were infected with *Lm*-OVA 1 day later. Naïve CD8^+^ or *Lm*-OVA-activated (Thy1.1^+^) OT-I cells were isolated from the spleen of infected mice at 3 dpi using an EasySep mouse CD90.1 (Thy1.1) positive selection kit (StemCell Technologies), as previously described^9,19^. For the *ex vivo* T cell killing assay, 1-5×10^5^ Thy1.1^+^CD8^+^ OT-I cells were transferred into Thy1.2^+^ C57BL/6J mice, followed by *Lm*-OVA infection 1 day later. At 7 dpi, *Lm*-OVA-activated (Thy1.1^+^) OT-I cells were isolated from the spleen of infected mice using an EasySep mouse CD90.1 (Thy1.1) positive selection kit (StemCell Technologies).

#### Flow cytometry

Single-cell suspensions were prepared by mechanical dissociation of harvested lymphoid tissues followed by red blood cell lysis, as previously described^9,19^. Single-cell suspensions (cultured cells or dissociated lymphoid tissues) were stained with a cocktail of fluorescently-labeled antibodies, lectins, and/or dyes listed in the **Key Resources Table**. Cell viability was assessed by using Fixable Viability Dye (FVD) eFluor 506 or FVD eFluor 780 according to the manufacturer’s protocols. Cell proliferation was assessed by Violet Proliferation Dye 450 (VPD450) dilution according to the manufacturer’s protocol. To assess cytokine production, splenocytes were isolated from *Lm*-OVA-infected mice at 7 dpi and stimulated with 1 μg/mL of OVA_257-264_ peptide and 50 U/mL IL-2 for 4 h, with GolgiStop (1:1500 dilution) added for the last 2 h of stimulation. After stimulation, cells were stained with surface marker antibodies in staining buffer (PBS with 2% FBS and 0.02% sodium azide) at 4°C for 1 h, fixed and permeabilized at 4°C for 1 h using the Foxp3/Transcription Factor Staining Buffer Set, and then stained with intracellular marker antibodies for either 1 h or overnight at 4°C. For cytokine profiling of Ametrine^+^ cells, cells were briefly fixed with 4% paraformaldehyde at room temperature for 5 min prior to fixation and permeabilization with the Foxp3/Transcription Factor Staining Buffer Set. For lectin staining, cells were stained with 2 μg/mL of each lectin individually at a density of 1×10^6^ cells/mL in staining buffer for 45 min at 4°C.

Analytical flow cytometry was performed on a Cytek Aurora or BD Accuri C6 Plus cytometer. Cell sorting was performed on an Astrios or BD FACSAria Fusion cytometer. Data analysis was performed using FlowJo (v9.9.5) software.

#### Extracellular flux analysis

T cell oxygen consumption rate (OCR) and extracellular acidification rate (ECAR) were measured by a Seahorse XF96 Extracellular Flux Analyzer as previously described^16^. Briefly, activated NTC- or sg*Ugcg*-modified CD8^+^ T cells (1×10^5^/well) were seeded in Seahorse XF medium containing 5 mM glucose, 0.5 mM glutamine, and 1 mM sodium pyruvate and centrifuged in a poly-D-lysine-coated XF96 plate. Cellular bioenergetics was assessed at regular time intervals following the sequential addition of oligomycin (2 μM), carbonyl cyanide 4-(trifluoromethoxy)phenylhydrazone (FCCP; 1.5 μM), rotenone/antimycin A (0.5 μM each), and monensin (20 μM). Data were normalized to cell number. Bioenergetics data analysis was based on protocols developed by Mookerjee et al.^70^, as previously described^16^.

#### Stable isotope labeling

Stable isotope labeling experiments in *in vitro*-activated T cells using liquid chromatography coupled to mass spectrometry (LC-MS) were conducted as previously described^15^. Briefly, CD8^+^ T cells were activated as above, washed with VIM medium, and re-seeded (e.g., 2 x 10^6^ cells/well in a 24-well plate for up to 2 h, or 2-3 x 10^6^ cells/well in a 6-well plate for 24 h) for the indicated times in VIM medium containing 10% dFBS, 50 μM 2-mercaptoethanol, and 5 mM [U-^13^C]-glucose. Mouse T cells were further supplemented with 50 U/mL of recombinant murine IL-2 and human T cells with 300 U/mL of recombinant human IL-2. Cells were transferred from tissue culture plates to tubes and centrifuged at 600 RCF for 2 min at 4°C. The cell pellet was washed with ice-cold saline before being snap-frozen on dry ice and stored at −80°C. For *ex vivo* stable isotope labeling experiments, Thy1.1^+^ OT-I cells were isolated from the spleen of *Lm*-OVA-infected mice at 3 dpi by magnetic bead sorting as previously described^9,19^, washed with VIM medium, and re-seeded at 2 x 10^6^ cells/well in a 24-well plate for the indicated times in VIM medium containing 10% dFBS, 50 μM 2-mercaptoethanol, 5 mM [U-^13^C]-glucose, and 50 U/mL of recombinant murine IL-2.

#### Stable isotope in vivo infusions

[U-^13^C]-glucose *in vivo* infusions of *Lm*-OVA-infected mice at 3 dpi were conducted as previously described^9,16,19^. Briefly, mice were fasted for 4 h prior to infusion. Mice were anesthetized using isoflurane and infused intravenously with 100 mg/mL [U-^13^C]-glucose (dissolved in sterile saline) over a 2 h period. An initial [U-^13^C]-glucose bolus of 0.6 mg/g mouse was administered, followed by a 2 h [U-^13^C]-glucose infusion at a rate of 0.125 μL/min/g mouse. Mice were euthanized via cervical dislocation and blood was harvested via cardiac puncture. Blood was centrifuged at 17,000 RCF for 10 min at 4°C to isolate serum, which was snap-frozen on dry ice and stored at −80°C. Spleens were harvested and processed for naïve CD8^+^ or Thy1.1^+^ T cell isolation as previously described^9,19^. Isolated T cells were snap-frozen on dry ice and stored at −80°C. T cells and serum were processed for metabolomics and lipidomics analyses as described below.

#### Metabolite and lipid extraction for LC-MS

Several extraction approaches were used depending on specific experimental goals. For experiments requiring both metabolite and lipid quantification, a Bligh-Dyer extraction^71^ (2:2:1.8 v/v chloroform:methanol:water) was used. Samples were homogenized in a 1:1 mixture of ice-cold chloroform (Sigma, 1024441000) and methanol (Fisher Scientific, A456). After solvent addition, extracts were vortexed for 10 s, sonicated for 5 min, and incubated on wet ice for 30 min. Incubation was followed by the addition of 0.9 volumes of LC-MS-grade water (Fisher Scientific, W6) to achieve the final 2:2:1.8 ratio. For serum samples, water content of the sample was accounted for (reported as 92%) to precisely maintain the 2:2:1.8 ratio when performing the water addition. Samples were then vortexed for an additional 10 s and followed with incubation on wet ice for 10 min.

For experiments requiring only metabolomics, a single phase AMW20 (40% acetonitrile, 40% methanol, and 20% water) extraction was used. For experiments requiring only lipidomics, a HEX:IPA (50% hexane and 50% isopropanol) extraction was used. For both AMW20 and HEX:IPA extracts, samples were homogenized with either ice-cold 4:4:2 acetonitrile:methanol:water or 1:1 hexane:isopropanol, respectively. After solvent addition, extracts were vortexed for 10 s, sonicated for 5 min, and incubated on wet ice for 1 h. Following the final incubation steps, Bligh-Dyer, AMW20, and HEX:IPA samples were centrifuged at 17,000 RCF for 10 min at 4°C. 32 μL serum equivalents of aqueous phase supernatant and 11.6 μL serum equivalents of organic phase subnatant were collected for metabolomics and lipidomics, respectively, and dried in a vacuum evaporator. In all extraction types, the extraction solvent:sample ratio was controlled within each experiment to avoid influence of varying extraction efficiency. However, for *in vivo* [U-^13^C]-glucose infusion experiments, T cells were extracted in a fixed 1 mL volume of chloroform:methanol:water (2:2:1.8), and protein content of the insoluble fraction was used to correct detected peak areas to starting cell amount.

Aqueous metabolomics samples were resuspended in LC-MS-grade water, and organic lipidomics samples were resuspended in a 1:1 mixture of LC-MS-grade isopropanol (Fisher Scientific, A461) and acetonitrile (Fisher Scientific, A955). 640 nL serum equivalents were injected on column for metabolomics, and 116 nL serum equivalents were injected on column for lipidomics. Injected on-column cell equivalents varied per experiment based on cell availability and ranged from 48,855-276,800 cell equivalents for metabolomics and 96,000-276,800 cell equivalents for lipidomics.

#### Metabolomics analysis

Metabolomic profiling and stable isotope tracing data were collected using a Vanquish LC system coupled to an Orbitrap Exploris 240 (Thermo Fisher Scientific) using a heated electrospray ionization (H-ESI) source in negative mode, as previously described^16,72^. 2 μL of each standard and/or sample was injected and run through a 24-min reversed-phase chromatography Zorbax RRHD extend-C18 Column (1.8 μm, 2.1 mm x 150 mm; Agilent, 759700-902) combined with a Zorbax extend-C18 guard column (1.8 μm, 2.1 mm x 5 mm; Agilent, 821725-907). Mobile phase A consisted of LC-MS-grade water with 3% LC-MS-grade methanol, mobile phase B was LC-MS-grade methanol and both mobile phases contained 10 mM tributylamine (Sigma, 90780), 15 mM LC-MS-grade acetic acid (Fisher Scientific, A11350), and 0.01% medronic acid (v/v; Agilent, 5191-4506). For the wash gradient, mobile phase A was kept the same, and mobile phase B was 99% LC-MS-grade acetonitrile. Column temperature was kept at 35°C, flow rate was held at 0.25 mL/min, and the chromatography gradient was as follows: 0-2.5 min held at 0% B, 2.5-7.5 min from 0% B to 20% B, 7.5-13 min from 20% B to 45% B, 13-20 min from 45% B to 99% B, and 20-24 min held at 99% B. A 16-min wash gradient was run in reverse flow direction between every injection to back-flush the column and to re-equilibrate solvent conditions as follows: 0-3 min held at 100% B and 0.25 mL/min, 3-3.5 min held at 100% B and ramp to 0.8 mL/min. 3.5-7.35 min held at 100% B and 0.8 mL/min, 7.35-7.5 min held at 100% B and ramp to 0.6 mL/min, 7.5-8.25 min from 100% B to 0% B and ramp to 0.4 mL/min, 8.25-15.5 min held at 0% B and ramp to 0.25 mL/min, and 15.5-16 min held at 0% B and 0.25 mL/min. Mass spectrometer parameters were: source voltage −2500 V, sheath gas 60, aux gas 19, sweep gas 1, ion transfer tube temperature 320°C, and vaporizer temperature 250°C. Full scan data were collected using the Orbitrap with a scan range of 70-850 m/z at a resolution of 240,000 and RF lens at 35%. Fragmentation was induced in the Orbitrap using assisted higher-energy collisional dissociation (HCD) collision energies at 15, 30, and 45%. Orbitrap resolution was 15,000, the isolation window was 2 m/z, and data-dependent scans were capped at 5 scans. Targeted mass, data-dependent MS2 (ddMS2) triggers were included for a panel of compounds (**Table S3**).

To chromatographically resolve UDP-Glc and UDP-Gal, we utilized a BEH amide chromatography with high-pH mobile phases on a Thermo Vanquish Horizon LC system coupled to an Orbitrap ID-X (Thermo Fisher Scientific) using an H-ESI source in negative mode. 2 μL of each standard and/or sample was injected on column. The 20-min chromatography used an Acquity UPLC BEH Amide Column (1.7 μm, 2.1 mm x 150 mm; Waters, 176001909) combined with an Acquity UPLC BEH Amide VanGuard Pre-column (1.7 μm, 2.1 mm x 5 mm; Waters, 186004799). Buffer A consisted of 100% LC-MS-grade water, 0.1% ammonium hydroxide (Fisher Scientific, A470), 0.1% medronic acid, 10 mM ammonium acetate (Sigma, 73594), and buffer B consisted of 90% LC-MS-grade acetonitrile, 10% LC-MS-grade water, 0.1% ammonium hydroxide, 0.1% medronic acid, and 10 mM ammonium acetate. Column temperature was kept at 40°C, flow rate was held at 0.4 mL/min, and the chromatography gradient was as follows: 0-1 min from 100% B to 90% B, 1-12.5 min from 90% B to 75% B, 12.5-19 min from 75% B to 60% B, and 19-20 min held at 60% B. A 30-min wash gradient was run between every injection to flush the column and to re-equilibrate solvent conditions as follows: 0-1 min held at 65% B and 0.4 mL/min, 1-13 min held at 65% B and ramp to 0.8 mL/min, 13-24 min held at 65% B and 0.8 mL/min, 24-24.5 min from 65% B to 100% B and held at 0.8 mL/min, 24.5-26 min held at 100% B and ramp to 1.2 mL/min, 26-28 min held at 100% B and 1.2 mL/min, 28-29 min held at 100% B and ramp to 0.4 mL/min, and 29-30 min held at 100% B and 0.4 mL/min. For Chromatography 1, the mass spectrometer parameters were: source voltage −2500 V, sheath gas 60, aux gas 19, sweep gas 1, ion transfer tube temperature 300°C, and vaporizer temperature 250°C. Full scan data were collected using the Orbitrap with a scan range of 70-1000 m/z at a resolution of 120,000 and RF lens at 35%. In tandem with the full scan, a targeted single ion scan (tSIM) was performed in the Orbitrap to target UDP-Glc and UDP-Gal and all their ^13^C isotopologues with a center mass of 565.0477 m/z and an isolation window 32 m/z. Resolution was set at 60,000, RF lens at 60%, and scan time was set for 10.5-13.5 min of the chromatographic gradient described above. ddMS2 fragmentation was induced in the Orbitrap using assisted HCD collision energies at 20, 40, 60, 80, 100% as well as with collision-induced dissociation (CID) at a collision energy of 35%. For both MS2 fragmentations, Orbitrap resolution was 30,000, the isolation window was 1.5 m/z, and total cycle time was 0.6 s. A targeted mass MS2 trigger for UDP-Hex (565.0477 m/z) was included.

Peak picking and integration were conducted in Skyline (v22-23) using in-house curated compound lists of accurate mass MS1 and retention time of chemical standards^72^. For tracing studies, this list was expanded to include all possible ^13^C isotopologues and natural abundance correction was completed using IsoCorrectR^73^.

#### Lipidomics analysis

Lipidomics analysis was completed with a Thermo Vanquish Horizon LC system coupled to an Orbitrap ID-X (Thermo Fisher Scientific) using an H-ESI source in positive mode^74^. Lipids were separated with a 30-min reversed-phase chromatography Accucore C30 column (2.6 μm, 2.1 mm x 150 mm; Thermo Fisher Scientific, 27826-152130) combined with an Accucore C30 guard column (2.6 μm, 2.1 mm x 10 mm; Thermo Fisher Scientific; 27826-012105). Buffer A consisted of 60% LC-MS-grade acetonitrile, 40% LC-MS-grade water, 0.1% LC-MS-grade formic acid (Fisher Scientific, A117), and 10 mM ammonium formate (Fisher Scientific, 70221), and buffer B consisted of 90% LC-MS-grade isopropanol, 8% LC-MS-grade acetonitrile, 2% LC-MS-grade water, 0.1% LC-MS-grade formic acid, and 10 mM ammonium formate. Column temperature was kept at 50°C, flow rate was held at 0.4 mL/min, and the chromatography gradient was as follows: 0-1 min held at 25% B, 1-3 min from 25% B to 40% B, 3-19 min from 40% B to 75% B, 19-20.5 min 75% B to 90% B, 20.5-28 min from 90% B to 95% B, 28-28.1 min from 95% B to 100% B, and 28.1-30 min held at 100% B. A 20-min wash gradient was run between every injection to flush the column and to re-equilibrate solvent conditions as follows: 0-10 min held at 100% B, 10-14 min from 100% B to 25% B, and 14-20 min held at 25% B. The mass spectrometer parameters were: source voltage 3250 V, sheath gas 40, aux gas 10, sweep gas 1, ion transfer tube temperature 300°C, and vaporizer temperature 275°C. Full scan data were collected using the Orbitrap with a scan range of 200-1700 m/z at a resolution of 500,000 and RF lens at 45%. ddMS2 fragmentation was induced in the Orbitrap using assisted HCD collision energies at 15, 30, 45, 75, 110% as well as with CID at a collision energy of 35%. For both MS2 fragmentations, Orbitrap resolution was 15,000 and the isolation window was 1.5 m/z. A m/z 184 mass trigger, indicative of phosphatidylcholines, was used for CID fragmentation. Data-dependent MS3 fragmentation was induced in the ion trap with scan rate set at “Rapid” using CID at a collision energy of 35%. MS3 scans were triggered by specific acyl chain losses for detailed analysis of mono-, di-, and triacylglycerides. Total cycle time was 2 s. Lipid identifications were assigned using LipidSearch (v5.0; Thermo Fisher Scientific), which was used to generate a compound list, and lipid peak areas were quantified in Skyline as described above.

#### Glycogen quantification

Cells were cultured in VIM medium containing 10% dFBS and 50 U/mL IL-2 for 24 h. 1×10^6^ cells were processed for glycogen quantification using a Glycogen-GLO Assay Kit (Promega) as per the manufacturer’s instructions. Luminescence was quantified 60 min after the addition of the glucose detection reagent using a BioTek Synergy Neo2 multi-mode plate reader. Glycogen abundance was normalized to protein content, which was measured using a Pierce Rapid Gold BCA Protein Assay Kit (Thermo Fisher Scientific).

#### ImmunobloĖng

Cells were lysed in RIPA buffer supplemented with protease and phosphatase inhibitor cocktails (Roche) on ice for ≥30 min. A Pierce BCA Protein Assay Kit (Thermo Fisher Scientific) was used to quantify protein from whole cell lysates. Equal amounts of protein were diluted in Laemmli sample buffer, boiled for 5 min, and resolved by SDS-PAGE on a 10% gel. Proteins were transferred onto nitrocellulose membranes. Membranes were blocked for 1 h in 5% non-fat milk in TBST at room temperature and incubated with primary antibodies against UGP2 or β-ACTIN (both 1:1000-diluted in 5% non-fat milk) overnight at 4°C. Membranes were washed three times for 5 min with 1x TBST, and then incubated for 1 h at room temperature with an HRP-conjugated secondary antibody diluted 1:2000 in 5% non-fat milk/TBST. Membranes were washed three times with 1x TBST and then developed using Enhanced Chemiluminescence (ECL) Western Blotting Detection Reagent (Cytivia). Antibodies are listed in the **Key Resources Table**.

#### Proteomics sample preparation and analysis

5×10^6^ control and UGCG-deficient CD8^+^ T cells were washed twice with ice-cold PBS and cell pellets were snap-frozen on dry ice. Cell lysates were extracted and digested using EasyPrep MS Sample Prep Kits (Thermo Fisher Scientific) according to the manufacturer’s instructions. Dried samples were resuspended in 0.1% trifluoroacetic acid.

For global proteome quantitation analysis, data-independent acquisition (DIA) analyses were performed on an Orbitrap Eclipse mass spectrometer coupled with the Vanquish Neo LC system (Thermo Fisher Scientific). The FAIMS Pro source was positioned between the nanoESI source and the mass spectrometer. A total of 2 μg of digested peptides were separated on a nano capillary column (20 cm x 75 μm I.D., 365 μm O.D., 1.7 μm C18; CoAnn Technologies) at a flow rate of 300 nL/min. Mobile phase A was water with 0.1% formic acid, and mobile phase B was 20% water and 80% acetonitrile with 0.1% formic acid. The LC gradient was programmed as follows: 1% B to 24% B over 110 min, 85% B over 5 min, and 98% B over 5 min, resulting in a total gradient length of 120 min. For FAIMS, selected compensation voltages (−40V, −55V, −70V) were applied throughout the LC-MS/MS runs. Full MS spectra were collected at a resolution of 120,000 (FWHM), and MS2 spectra at 30,000 (FWHM). Standard automatic gain control (AGC) targets and automatic maximum injection times were used. A precursor range of 380-980 m/z was set for MS2 scans, with a 50 m/z isolation window and a 1 m/z overlap for each scan cycle. A 32% HCD collision energy was used for MS2 fragmentation.

To generate a hybrid library for directDIA analysis in Spectronaut, pooled samples underwent data-dependent acquisition (DDA) employing 11 distinct FAIMS CV settings ranging from −30 to 80 CV. Full MS spectra were collected at 120,000 resolution (FWHM), and MS2 spectra at 30,000 resolutions (FWHM). The standard AGC target and automatic maximum injection time were selected. Ions with charges of 2-5 were filtered. An isolation window of 1.6 m/z was used with quadrupole isolation mode, and ions were fragmented using HCD with a collision energy of 32%.

For the targeted quantitation, parallel reaction monitoring (PRM) was performed on an Exploris 480 mass spectrometer coupled with the Vanquish Neo LC system (Thermo Fisher Scientific). A total of 2 μg of digested peptides were separated on a nano capillary column (20 cm x 75 μm I.D., 365 μm O.D., 1.7 μm C18; CoAnn Technologies) at a flow rate of 300 nL/min. Mobile phase A was water with 0.1% formic acid, and mobile phase B was 20% water and 80% acetonitrile with 0.1% formic acid. The LC gradient was as follows: 1% B to 26% B over 51 min, 85% B over 5 min, and 98% B over 4 min, with a total gradient length of 60 min. Full MS spectra (m/z 375-1200) were collected at a resolution of 120,000 (FWHM), and MS2 spectra at 30,000 (FWHM). Standard AGC targets and automatic maximum injection times were used for both full and MS2 scans. A 32% HCD collision energy was used for MS2 fragmentation. All samples were analyzed using a multiplexed PRM method with a scheduled inclusion list containing the target precursor ions. Three unique peptides of UGCG (QGFAATLEQVYFGTSHPR, VGLVHGLPYVADR, and SYISANVTGFK) were used to measure the relative abundance of UGCG, while two α-ACTININ 4 (ACTN4) peptides (DDPVTNLNNAFEVAEK and LVSIGAEEIVDGNAK) served as internal standards. Synthetic peptides were obtained from Biosynth.

DIA data were processed in Spectronaut v18 (Biognosys) using directDIA analysis. Data were searched against the *Mus musculus* reference proteome database (Uniprot, Taxon ID: 10090) with the manufacturer’s default parameters. Briefly, trypsin/P was set as the digestion enzyme, allowing for two missed cleavages. Cysteine carbamidomethylation was set as a fixed modification, while methionine oxidation and protein N-terminus acetylation were set as variable modifications. Identification was performed using a 1% q value cutoff at both the precursor and protein levels. Both peptide precursor and protein false discovery rates were controlled at 1%. Ion chromatograms of fragment ions were used for quantification, with the area under the curve between the XIC peak boundaries calculated for each targeted ion. DDA raw files were utilized in Library Extension Runs to enhance proteome coverage to generate a hybrid library. All PRM data analysis and integration were performed using Skyline software. The transitions’ intensity rank order and chromatographic elution were required to match those of a synthetic standard for each measured peptide.

Differential abundance of proteins was analyzed similarly to House *et al.*^72^ using the R v4.4.0 package limma (v3.60.0). Briefly, LIMMA-eBayes was fit on all proteins with less than 30% missingness and any remaining missing values were imputed using left-censored methods from imputeLCMD (v2.1). Second generation p values were then calculated using a null interval of 0.75-1.25 fold-differences to reduce the Type 1 Error rate and improved interpretability^75^. Gene set enrichment analysis (GSEA) on differentially abundant proteins was conducted using the gseGO function via clusterProfiler^76^ (v4.12.0) and examined all of: biological process, cellular component, and molecular function. Significance of enrichment was based on Benjamini-Hochberg-adjusted p values^77^. Results of gseGO were visualized through network plots via the aPEAR package^78^ (v1.0.0) and distinct pathway clusters from this analysis were used to calculate average normalized enrichment scores to represent overall enrichment in activated and suppressed pathways. The pipelines used for this analysis are hosted on GitHub: https://github.com/vari-bbc/Ugcg-KO_proteomics_analysis.

#### T cell killing assay

C57BL/6J mice were adoptively transferred with 1-5×10^5^ Thy1.1^+^CD8^+^ OT-I cells (NTC- or sg*Ugcg*-modified) via the tail vein and infected with *Lm*-OVA the following day. At 7 dpi, Thy1.1^+^ cells were rapidly isolated from the spleen using magnetic beads via positive selection. tdTomato-expressing MC38-OVA cells^16^ were seeded in a 96-well plate (5×10^3^ cells/well). 24 h later, a range of 2×10^5^ to ∼1563 *Lm*-OVA-activated NTC- or sg*Ugcg*-modified CD8^+^ OT-I cells were added to the MC38-OVA cells in standard IMDM supplemented with 50 U/mL IL-2. tdTomato^+^ cells were quantified over a 24 h period using IncuCyte live-cell analysis (Sartorius). The percentage of dead MC38-OVA tumor cells was calculated by normalizing the number of tdTomato-expressing cells to media-only (no T cell) controls.

#### B16-OVA tumor model and OT-I cell adoptive transfer

Female C57BL/6J mice were injected with 5×10^5^ B16-OVA cells subcutaneously in the abdominal flank. Once palpable tumors were present, tumor measurements were obtained every 2-3 days using a caliper. 7 days after B16-OVA cell injection, mice received 1×10^6^ Thy1.1^+^CD8^+^ OT-I cells (NTC- or sg*Ugcg*-modified) or HBSS as a vehicle control intravenously via the tail vein. Mice were euthanized as they reached human endpoints, which included a maximum tumor volume of ≥1500 mm^3^. For tumor-infiltrating lymphocyte (TIL) analysis, tumors were harvested 14 days after B16-OVA cell injection (7 days after OT-I adoptive transfer) and mechanically disrupted to generate a single-cell suspension. Cells were pelleted (500 RCF for 5 min at 4°C) and resuspended in red blood cell lysis buffer (0.1 mM EDTA, 155 mM NH_4_Cl, and 12 mM NaHCO_3_) for 1 min at room temperature. Cells were stained and analyzed by flow cytometry as described in the “*Flow cytometry*” section.

#### TCR crosslinking and lipid raft aggregation analysis

Activated NTC- or sg*Ugcg*-modified CD8^+^ T cells were starved of IL-2 for 4 hr. Cells were incubated with 10 μg/mL anti-CD3ε (clone 145-2C11) on ice for 30 min, and then with 10 μg/mL biotin-conjugated goat anti-Armenian hamster IgG at 37°C for 30 min. Sodium azide was added at a final concentration of 0.2% w/v to terminate the reaction. 10,000 cells in 100 μL of PBS were transferred to 13 mm single ring slides (Fisherbrand) using a Shandon Cytospin 3 centrifuge (1,000 RPM for 5 min). Cells were then fixed in 10% neutral-buffered formalin for 10 min at room temperature, washed twice with PBS (5 min each), and washed once with reverse osmosis (RO) water for 5 min. Slides were allowed to dry overnight at room temperature and then stored at 4°C until ready to stain.

Prior to staining, cells were washed twice with PBS (5 min each) and blocked with 2% FBS for 30 min at room temperature. Antibody/stain dilutions were performed using Antibody Diluent (Dako): CTxB-FITC (1:100) and streptavidin-Alexa Fluor 647 (1:500). Cells were stained overnight at 4°C. Cells were counterstained with DAPI for 10 min at room temperature prior to cover slip mounting with ProLonged Gold Antifade Mountant (Thermo Fisher). Slides were stored at 4°C (protected from light) until ready to image.

Confocal Z-stacks were collected using a Zeiss LSM 880 equipped with an Axio Observer 7 inverted microscope body and acquired with Zen Black (v2.3) software using 405 nm diode, 488 nm argon ion, and 633 nm HeNe laser lines, plus a transmitted light channel and a 0.92 AU pinhole. Emitted light was detected through a Zeiss Plan-apochromat 63x/1.4 NA oil immersion objective, using a PMT GaAsP detector. Images were collected sequentially at 1024×1024 pixel resolution, using 0.5 μm z-steps. Individual voxels were therefore 0.13 μm x 0.13 μm x 0.5 μm (xyz). A pixel dwell time of 0.42 μs and an optical zoom of 1.0 were used for the collection of all images. Large fields of view were obtained by automated collection and stitching of a 3×3 rectangular grid with 10% overlap.

Confocal Z-stacks were converted to max projections, channels split and saved as a TIFF using Fiji v2.15.1 prior to importing into CellProfiler for analysis^79,80^. The maximum projections for the DAPI, anti-streptavidin (CD3ε), and CTxB channels were used to quantify the number, location, and fluorescence distribution of the TCRs and lipid rafts per cell. Briefly, nuclei and their approximate cytoplasmic outlines were generated with the IdentifyPrimaryObjects and IdentifySecondaryObjects modules using the DAPI and CTxB channels, respectively. The objects from IdentifySecondaryObjects module were then used to mask the TCR channel; IdentifyPrimaryObjects module was used to segment the TCR puncta in this masked channel. Once all objects were generated and related to each other, a series of measurement modules were used to quantify fluorescence intensity, distribution, and texture, and shape and size of detected puncta. The CellProfiler pipeline used for this analysis is hosted on GitHub: https://github.com/vaioic/Published_Scripts. All measurements were exported to a CSV file for further statistical analysis. Representative images were generated using the QuickFigures Fiji plugin^81^.

### QUANTIFICATION AND STATISTICAL ANALYSIS

Unless otherwise stated, data are presented as mean ± standard deviation (SD) for technical replicates and mean ± standard error of the mean (SEM) for biological replicates. Observations from technical replicate data were reproduced in at least two independent experiments. Statistical analyses were performed using GraphPad Prism software (GraphPad). A two-tailed, unpaired Student’s t-test was used to compare the means between two independent groups. All other statistical tests are specified in the corresponding figure legends. Statistical significance is indicated in all figures by the following annotations: *, p < 0.05; **, p < 0.01; ***, p < 0.001; ****, p < 0.0001; ns, not significant.

## SUPPLEMENTAL INFORMATION

**Document S1** (.pdf) – Figures S1-S6.

**Table S1** (.csv) – Differentially expressed proteins in control and UGCG-deficient CD8^+^ T cells, related to Figure 6.

**Table S2** (.csv) – gseGO pathway analysis, related to Figure 6.

**Table S3** (.xlsx) – ddMS2 inclusion list for ion paired metabolomics.

