## Supplemental Figures S1-S6 for "Glucose-dependent glycosphingolipid biosynthesis fuels CD8^+^ T cell function and tumor control"

**Figure S1**

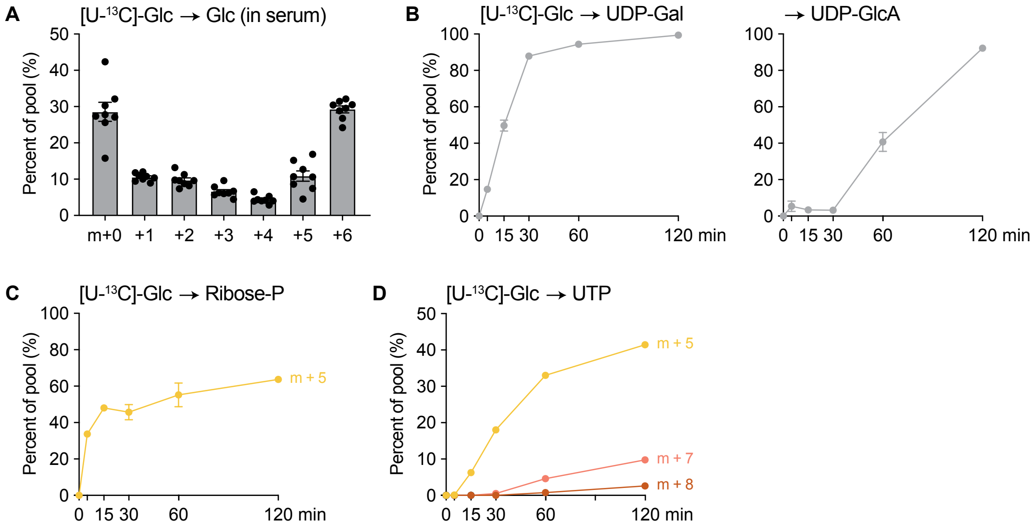


**Figure S1, related to Figure 1. [U-$^{13}$C]-glucose tracing in *Lm*-OVA-activated CD8$^{+}$ T cells *in vivo* and *ex vivo*.** (**A**) Mass isotopologue distribution (MID) of glucose in serum following 2 h of [U-$^{13}$C]-glucose infusion *in vivo* in *Lm*-OVA-infected mice at 3 dpi. Data represent the mean ± SEM (n = 8 mice/group). (**B-D**) CD8$^{+}$ OT-I Teff cells isolated from *Lm*-OVA-infected mice at 3 dpi were cultured *ex vivo* for up to 2 h in VIM medium containing 5 mM [U-$^{13}$C]-glucose. (**B**) Timecourse of [U-$^{13}$C]-glucose incorporation into UDP-galactose (UDP-Gal; *left*) and UDP-glucuronic acid (UDP-GlcA; *right*). Total $^{13}$C enrichment (% of pool) from [U-$^{13}$C]-glucose is shown. Data represent the mean ± SEM (n = 3 biological replicates). (**C**) [U-$^{13}$C]-glucose-derived m+5 ribose-P in CD8$^{+}$ Teff cells over time. Data represent the mean ± SEM (n = 3 biological replicates). (**D**) MID of [U-$^{13}$C]-glucose-derived UTP in CD8$^{+}$ Teff cells over time. Data represent the mean ± SEM (n = 3 biological replicates).

#### Figure S2

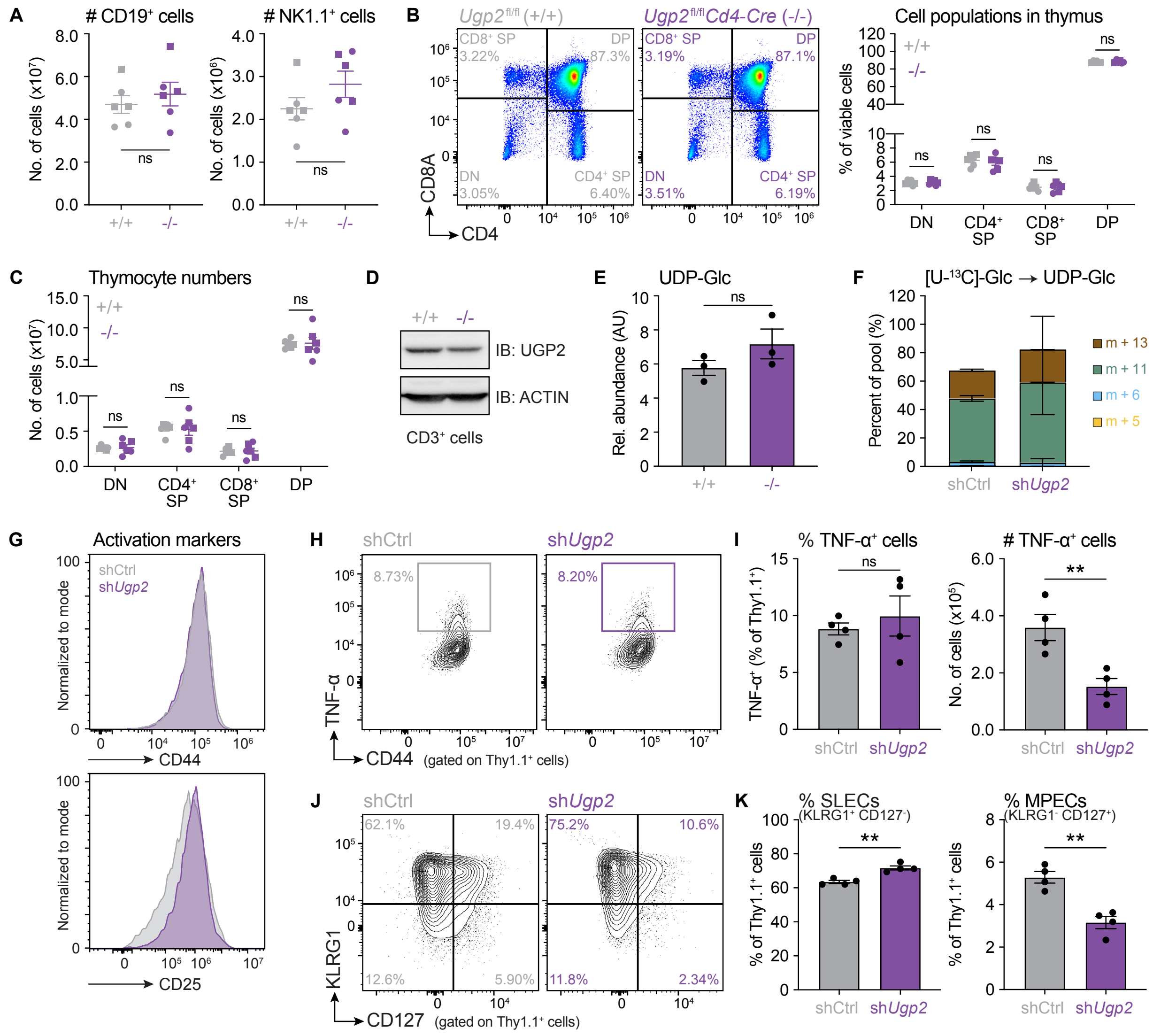

A
### CD19+ cells
### NK1.1+ cells
No. of cells (x10^7)
No. of cells (x10^6)
ns
+/+
-/-
B
Ugp2fl/fl (+/+)
Ugp2fl/fl Cd4-Cre (-/-)
CD8+ SP
DP
DN
CD4+ SP
3.22%
87.3%
3.05%
6.40%
3.19%
87.1%
3.51%
6.19%
CD8A
CD4
Cell populations in thymus
% of viable cells
DN
CD4+ SP
CD8+ SP
DP
C
Thymocyte numbers
No. of cells (x10^7)
D
IB: UGP2
IB: ACTIN
CD3+ cells
E
UDP-Glc
Rel. abundance (AU)
F
[U-13C]-Glc → UDP-Glc
Percent of pool (%)
m + 13
m + 11
m + 6
m + 5
shCtrl
shUgp2
G
Activation markers
Normalized to mode
CD44
CD25
H
8.73%
8.20%
TNF-α
CD44 (gated on Thy1.1+ cells)
I
% TNF-α+ cells
### TNF-α+ cells
TNF-α+ (% of Thy1.1+)
No. of cells (x10^5)
**
J
62.1%
19.4%
12.6%
5.90%
75.2%
10.6%
11.8%
2.34%
KLRG1
CD127 (gated on Thy1.1+ cells)
K
% SLECs (KLRG1+ CD127-)
% MPECs (KLRG1- CD127+)
% of Thy1.1+ cells

**Figure S2, related to Figure 2. Characterization of the *Ugp2*$^{fl/fl}$*Cd4-Cre* mouse model and responses of sh*Ugp2*-expressing CD8$^{+}$ T cells to *Lm*-OVA infection *in vivo*.** (**A**) Number of splenic CD19$^{+}$ B cells (*left*) and NK1.1$^{+}$ NK cells (*right*) in *Ugp2*$^{fl/fl}$ (+/+) and *Ugp2*$^{fl/fl}$*Cd4-Cre* (-/-) mice. Data represent the mean ± SEM (n = 6 mice/group; circle = female, square = male). (**B**) *Left*, Representative flow cytometry plots for CD8A versus CD4 expression on thymocytes isolated from *Ugp2*$^{fl/fl}$ (+/+) and *Ugp2*$^{fl/fl}$*Cd4-Cre* (-/-) mice. *Right*, Percent distribution of double-negative (DN), CD4 single-positive (CD4$^{+}$ SP), CD8 single-positive (CD8$^{+}$ SP), and double-positive (DP) cells in the thymus of *Ugp2*$^{fl/fl}$ (+/+) and *Ugp2*$^{fl/fl}$*Cd4-Cre* (-/-) mice. Data represent the mean ± SEM (n = 6 mice/group; circle = female, square = male). (**C**) Total number of DN, CD4$^{+}$ SP, CD8$^{+}$ SP, and DP cells in the thymus of *Ugp2*$^{fl/fl}$ (+/+) and *Ugp2*$^{fl/fl}$*Cd4-Cre* (-/-) mice. Data represent the mean ± SEM (n = 6 mice/group; circle = female, square = male). (**D**) Immunoblot of UGP2 protein expression in CD3$^{+}$ T cells isolated from *Ugp2*$^{fl/fl}$ (+/+) and *Ugp2*$^{fl/fl}$*Cd4-Cre* (-/-) mice. ACTIN protein expression is shown as a loading control. (**E**) Relative UDP-Glc abundance in purified CD3$^{+}$ T cells isolated from *Ugp2*$^{fl/fl}$ (+/+) and *Ugp2*$^{fl/fl}$*Cd4-Cre* (-/-) mice. Data represent the mean ± SEM (n = 3 mice/group). AU, arbitrary unit. (**F**) Mass isotopologue distribution (MID) of [U-$^{13}$C]-glucose-derived UDP-Glc in control (shCtrl) and sh*Ugp2*-expressing CD8$^{+}$ T cells cultured *in vitro* for 24 h in VIM medium containing 5 mM [U-$^{13}$C]-glucose. Data represent the mean ± SEM (n = 3). (**G**) Histograms of CD44 (*top*) and CD25 (*bottom*) expression at the surface of control (shCtrl) and sh*Ugp2*-expressing CD8$^{+}$ T cells cultured *in vitro*. (**H-I**) Cytokine response of UGP2-depleted CD8$^{+}$ T cells *ex vivo*. (**H**) Representative flow cytometry plots showing the percentage of TNF-$\alpha$-producing Thy1.1$^{+}$ control (shCtrl) and sh*Ugp2*-expressing CD8$^{+}$ OT-I cells in the spleen of *Lm*-OVA-infected mice at 7 dpi after *ex vivo* re-stimulation with OVA peptide. (**I**) Percentage (*left*) and total number (*right*) of TNF-$\alpha$-producing Thy1.1$^{+}$ control and sh*Ugp2*-expressing CD8$^{+}$ OT-I cells in the spleen of *Lm*-OVA-infected mice at 7 dpi after *ex vivo* re-stimulation with OVA peptide. Data represent the mean ± SEM (n = 4 mice/group). (**J-K**) Distribution of control and sh*Ugp2*-expressing CD8$^{+}$ OT-I cells in various differentiation states isolated from the spleen of *Lm*-OVA-infected mice at 7 dpi. (**J**) Representative flow cytometry plots of KLRG1 and CD127 expression on Thy1.1$^{+}$ control (shCtrl) and sh*Ugp2*-expressing CD8$^{+}$ OT-I cells. (**K**) Percentage of KLRG1$^{+}$CD127$^{-}$ short-lived effector cells (SLECs; *left*) and KLRG1$^{-}$CD127$^{+}$ memory precursor effector cells (MPECs; *right*) (of Thy1.1$^{+}$ cells). Data represent the mean ± SEM (n = 4 mice/group).

#### Figure S3

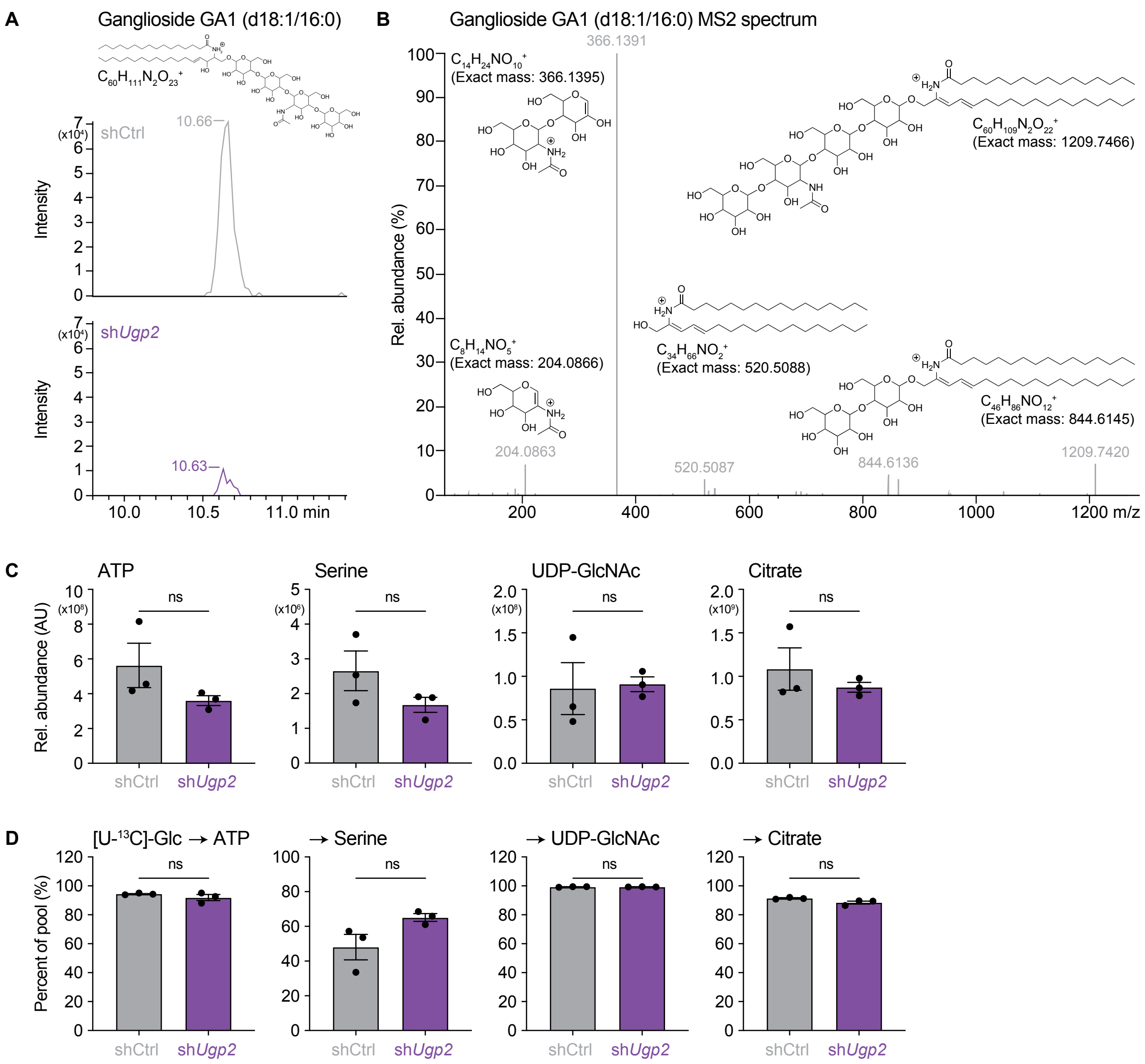


**Figure S3, related to Figure 3. GA1 ganglioside quantification and [U-$^{13}$C]-glucose tracing in control and UGP2-depleted CD8$^+$ T cells.** (**A-B**) Quantitation and identification of GA1 (d18:1/16:0) ganglioside by LC-MS. (**A**) Representative extracted ion chromatograms of m/z 1227.7572 (M+H, ± 5 ppm), which corresponds to the protonated chemical formula $C_{60}H_{111}N_2O_{23}$ of GA1 (d18:1/16:0), in control (shCtrl) and sh*Ugp2*-expressing CD8$^+$ T cells. (**B**) LC-MS/MS fragmentation analysis with inset fragment structures. (**C-D**) Control (shCtrl) and sh*Ugp2*-expressing CD8$^+$ T cells were cultured in VIM medium containing 5 mM [U-$^{13}$C]-glucose for 24 h prior to metabolite extraction. (**C**) Relative abundance of, or (**D**) glucose-derived $^{13}$C incorporation into, ATP, serine, UDP-GlcNAc, and citrate. Data represent the mean ± SEM (n = 3).

**Figure S4**

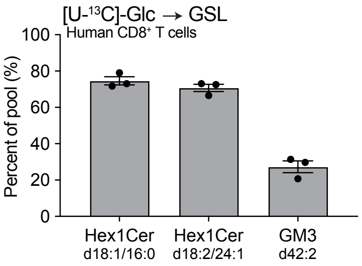


**Figure S4, related to Figure 4. Activated human CD8$^+$ T cells use glucose for GSL biosynthesis.** Human CD8$^+$ T cells were activated *in vitro* with anti-CD3 and anti-CD28 antibodies, and then cultured in VIM medium containing 5 mM [U-$^{13}$C]-glucose for 24 h prior to metabolite extraction. Glucose-derived $^{13}$C incorporation into hexosylceramide (Hex1Cer) species and GM3 (d42:2) ganglioside are shown. Data represent the mean ± SEM (n = 3 human donors).

#### Figure S5

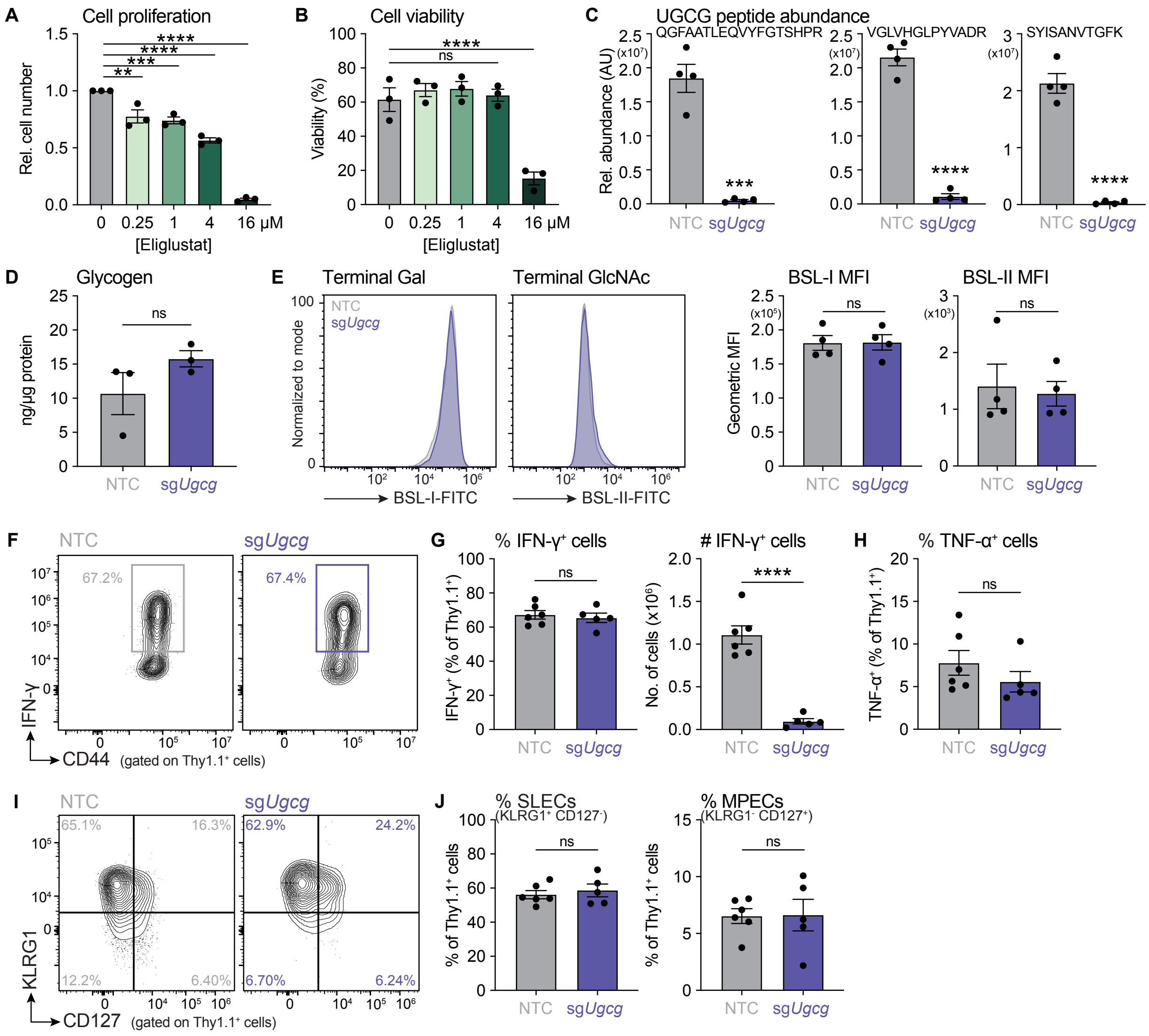

A
Cell proliferation
Rel. cell number
[Eliglustat]
0 0.25 1 4 16 µM
B
Cell viability
Viability (%)
C
UGCG peptide abundance
QGFAATLEQVYFGTSHPR
VGLVHGLPYVADR
SYISANVTGFK
Rel. abundance (AU)
NTC sgUgcg
D
Glycogen
ng/µg protein
E
Terminal Gal
Terminal GlcNAc
Normalized to mode
BSL-I-FITC
BSL-II-FITC
BSL-I MFI
BSL-II MFI
Geometric MFI
F
IFN-γ
CD44 (gated on Thy1.1⁺ cells)
67.2%
67.4%
G
% IFN-γ⁺ cells
### IFN-γ⁺ cells
IFN-γ⁺ (% of Thy1.1⁺)
No. of cells (x10⁶)
H
% TNF-α⁺ cells
TNF-α⁺ (% of Thy1.1⁺)
I
KLRG1
CD127 (gated on Thy1.1⁺ cells)
65.1% 16.3% 12.2% 6.40%
62.9% 24.2% 6.70% 6.24%
J
% SLECs (KLRG1⁺ CD127⁻)
% MPECs (KLRG1⁻ CD127⁺)
% of Thy1.1⁺ cells
ns

**Figure S5, related to Figure 5. Inhibiting GSL biosynthesis in CD8$^+$ T cells via eliglustat treatment or CRISPR/Cas9-mediated deletion of *Ugcg*.** (**A-B**) CD8$^+$ T cells were first activated with anti-CD3 and anti-CD28 antibodies *in vitro* and then treated with the indicated concentrations of eliglustat for 72 h. (**A**) Relative cell number and (**B**) cell viability of activated CD8$^+$ T cells after 72 h of treatment with eliglustat. Data represent the mean ± SEM (n = 3). Statistical significance was determined using a one-way ANOVA and Dunnett's multiple comparisons test. (**C**) Activated CD8$^+$ T cells were modified using CRISPR/Cas9 gene editing with either a non-targeting control (NTC) single guide RNA (sgRNA) or a sgRNA targeting *Ugcg* (sg*Ugcg*). Relative abundance of three different UGCG peptides in CD8$^+$ T cells as determined by LC-MS-based proteomics. Data represent the mean ± SEM (n = 4). (**D**) Total glycogen abundance in NTC- and sg*Ugcg*-modified CD8$^+$ T cells. Data represent the mean ± SEM (n = 3). (**E**) Representative histograms (*left*) and quantification of geometric mean fluorescence intensity (MFI; *right*) of BSL-I and BSL-II lectins on NTC- and sg*Ugcg*-modified CD8$^+$ T cells. Data represent the mean ± SEM (n = 4). (**F-H**) Cytokine response of UGCG-deficient CD8$^+$ T cells *ex vivo*. (**H**) Representative flow cytometry plots showing the percentage of IFN-$\gamma$-producing Thy1.1$^+$ NTC- (control) and sg*Ugcg*-modified CD8$^+$ OT-I cells in the spleen of *Lm*-OVA-infected mice at 7 dpi after *ex vivo* re-stimulation with OVA peptide. (**G**) Percentage (*left*) and total number (*right*) of IFN-$\gamma$-producing Thy1.1$^+$ control and sg*Ugcg*-modified CD8$^+$ OT-I cells in the spleen of *Lm*-OVA-infected mice at 7 dpi after *ex vivo* re-stimulation with OVA peptide. Data represent the mean ± SEM (n = 5-6 mice/group). (**H**) Percentage of TNF-$\alpha$-producing Thy1.1$^+$ control and sg*Ugcg*-modified CD8$^+$ OT-I cells in the spleen of *Lm*-OVA-infected mice at 7 dpi after *ex vivo* re-stimulation with OVA peptide. Data represent the mean ± SEM (n = 5-6 mice/group). (**I-J**) Distribution of control and sg*Ugcg*-modified CD8$^+$ OT-I cells in various differentiation states isolated from the spleen of *Lm*-OVA-infected mice at 7 dpi. (**I**) Representative flow cytometry plots of KLRG1 and CD127 expression on Thy1.1$^+$ control (NTC) and sg*Ugcg*-modified CD8$^+$ OT-I cells. (**K**) Percentage of KLRG1$^+$CD127$^-$ short-lived effector cells (SLECs; *left*) and KLRG1$^-$CD127$^+$ memory precursor effector cells (MPECs; *right*) (of Thy1.1$^+$ cells). Data represent the mean ± SEM (n = 5-6 mice/group).

**Figure S6**

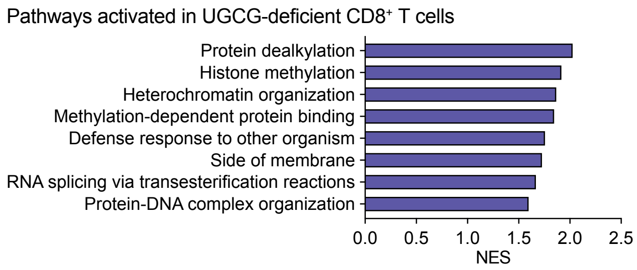


**Figure S6, related to Figure 6. Pathway analysis of differentially expressed proteins in control and UGCG-deficient CD8$^+$ T cells.** Pathway analysis indicating the top activated pathways in UGCG-deficient CD8$^+$ T cells from Figure 6A. NES, normalized enrichment score.
